# Cell division timing shapes the morphology and size of nascent multicellular organisms

**DOI:** 10.1101/2025.04.23.650085

**Authors:** Luis F. Cedeño-Pérez, Rozenn M. Pineau, Thomas C. Day, William C. Ratcliff, Peter L. Conlin

## Abstract

Upon making the transition from unicellularity to multicellularity, many previously optimized cellular traits experience the renewed scrutiny of natural selection due to their novel effects on multicellular phenotypes. However, we lack a comprehensive understanding of how and why specific cellular traits influence multicellular phenotypes and fitness. The snowflake yeast model system provides a tractable entry point for such investigations because cluster size is governed by simple topological and mechanical constraints and the connections between some cell-level traits (cellular aspect ratio, cell volume, bud neck strength) and group-level traits (group size) have been well characterized. However, differences in these known properties are insufficient to explain the difference in cluster size between the two strains that serve as the ancestors of the ongoing Multicellularity Long-Term Evolution Experiment (MuLTEE). Using time-lapse microscopy and single cell tracking, we identify cell division timing as a cellular trait that strongly influences multicellular morphology and size in snowflake yeast. The “petite” snowflake yeast ancestor divides asynchronously, with a 28% longer first division, while the “grande” ancestor divides synchronously. Using network theoretical and biophysical models, we show that strains exhibiting a first division delay generate more highly-branched network topologies, accelerating the accumulation of crowding-induced mechanical stress, resulting in clusters that fracture at smaller sizes. Conversely, synchronously dividing strains produce more symmetric, larger clusters. Synchronous cell division can provide benefits through both faster growth and larger size, suggesting multiple potential selective pathways for its evolution. Furthermore, we explore how accelerated first division can produce even larger groups and how another unexpected mechanism for modifying cluster topology, apoptosis rate, can exert a similar effect. Our results identify cell division timing as a previously underappreciated axis of phenotypic variation that strongly influences multicellular morphology and reproduction in the snowflake yeast system. This suggests that temporal regulation of cell division may be an evolutionarily accessible mechanism for early control of morphogenesis in nascent multicellular organisms with permanent intercellular bonds.

## Introduction

The evolution of multicellularity is one of life’s major evolutionary transitions, in which formerly free-living, lower-level units come together to form a new, higher-level entity (1–3). The first step in this transition is generally believed to be the formation of small, undifferentiated groups of cells (4,5). Once this new level of hierarchical organization has formed, if membership in the multicellular group affects a cell’s probability of reproduction or survival, the group’s existence has the potential to alter the relationship between cellular traits and fitness (6,7). These shifts in cellular trait optima are ultimately responsible for the opening of previously inaccessible evolutionary paths to new multicellular phenotypes, expansion into new niche space, and the adaptive radiations associated with the evolution of multicellularity in several clades (8). But what types of cellular traits have meaningful fitness consequences in the context of small, undifferentiated groups of cells? How does more sophisticated control over multicellular morphogenesis get off the ground?

All aspects of multicellular morphology and behavior must necessarily arise from the underlying properties and interactions of the component cells. There exists robust theory explaining how cellular traits can shape certain multicellular phenotypes: properties such as cellular growth rate, adhesion, polarity, and division patterning can strongly influence group morphology, mechanical stability, and life cycles (9,10). Moreover, many empirical studies have demonstrated tight links between specific cellular traits and group phenotypes. In the choanoflagellate *Choanoeca flexa*, for example, synchronized shape changes in individual cells drive collective sheet bending through direct microvilli tip adhesion (11,12). Changes in cell size and division number likewise underlie transitions from unicellular ancestors to multicellular groups in the volvocine algae (13–15), and in filamentous cyanobacteria, the spatial arrangement of dividing cells determines filament integrity and the emergence of specialized cell types (16–18). In *Dictyostelium discoideum*, which forms multicellular bodies by aggregation of free-living amoebae, heritable variation in cell-fate bias and adhesion shapes the geometry and robustness of multicellular fruiting bodies, with direct consequences for fitness (19–21). Each of these systems provides an example of a direct causal link between a cellular trait and a multicellular phenotype, yet we are far from a comprehensive, cell-level understanding of multicellular morphogenesis in even the simplest multicellular systems. To test how cellular traits give rise to multicellular phenotypes and morphogenesis, some researchers have turned to experimental evolution, using nascent multicellular organisms as tractable systems for directly observing the evolution of novel cellular traits and their consequences on group-level morphology and fitness.

Our work takes inspiration from the “snowflake yeast” system, a mutant derivative of the yeast *Saccharomyces cerevisiae* that forms undifferentiated multicellular clusters with permanent intercellular bonds via incomplete cell separation after cell division. This simple form of multicellularity evolved from a unicellular ancestor in 10/10 replicate populations in a matter of weeks in response to daily selection for rapid sedimentation in liquid media (22). Snowflake yeast have a simple form of group-level reproduction whereby single clusters will undergo fragmentation into two independently viable clusters via the fracture of a single mother-daughter cell-cell connection due to crowding-induced mechanical stress (23). Fragmentation due to the accumulation of mechanical stress, an inevitable consequence of growth for organisms with a branching tree architecture and permanent intercellular bonds, has been shown to be one of the primary barriers to evolving large group size. Accordingly, several key adaptations during the first 5,000 generations of the MuLTEE successfully prevent crowding (the evolution of increased cellular aspect ratio) or mitigate the effects of mechanical stress (the evolution of thicker, stronger cell-cell connections) (24,25). Interestingly, these known characteristics that increase group size only modify the biophysical properties of the group, such as its packing density and therefore its rate of accumulation of mechanical stress. In other words, they do not modify the underlying topological structure that connects cells. Another way to mitigate the effects of crowding-induced mechanical stress would be to change the topology of cell-cell connections; this possibility has not yet been investigated (though see (Libby et al., 2014; Nanda et al., 2024) for prior network modeling work that investigates how cellular packing density constrains the growth rate and size of multicellular yeast clusters).

Here we show that temporal regulation of cell division timing is an evolutionarily accessible mechanism that can modify a group’s underlying topology, providing early control over morphogenesis in nascent multicellular organisms with permanent intercellular bonds. We used soft-agar time-lapse microscopy and single-cell lineage tracing to collect detailed phenotypic measurements of individual cells during the growth of multicellular snowflake yeast clusters. We applied these methods to two simple, undifferentiated multicellular strains and discovered a previously unappreciated difference in the synchrony/asynchrony of cell division between the two strains. Through computational simulations and network modeling, we demonstrate that asynchrony driven by a first division delay, but not asynchrony driven by increased variation in doubling time, results in altered cluster topology and smaller fragmentation sizes compared to synchronously dividing strains, and that these morphological differences carry significant fitness consequences. Consistent with these predictions, clusters that began with asynchronous divisions rapidly evolved synchrony and maintained it for thousands of generations using strains from the ongoing Multicellularity Long-Term Evolution Experiment (MuLTEE) (24–27). We then explore how different patterns of asynchrony affect group properties, revealing that when cells accelerate their first division, they generate even larger clusters than synchronously dividing strains. Furthermore, we show that apoptosis offers a second independent lever for reshaping network topology, as apoptotic cells which stop dividing early can increase the size at fracture of the multicellular group. Additionally, we found that apoptosis interacts with cell division timing, reducing the potential benefits of an accelerated first cell division. Together, these results illustrate how modifications to cellular behavior reshape network topology, generating heritable variation in multicellular phenotypes upon which selection can act.

## Results

### Motivating observation

The two strains that function as the ancestors of the ongoing Multicellularity Long-Term Evolution Experiment (MuLTEE) genetically differ only in the presence or absence of a complete, functional mitochondrial genome (26). Both strains, which we refer to respectively as “grande” (respiration-competent) and “petite” (respiration-deficient), are homozygous diploid derivatives of Saccharomyces cerevisiae strain Y55. Multicellularity was engineered by homozygous replacement of the ACE2 transcription factor with a kanamycin resistance marker (ace2::KANMX/ace2::KANMX), resulting in the formation of small, undifferentiated multicellular clusters. The petite strain was then generated by selecting for a spontaneous mutant incapable of respiration (Figure 1A).

**Figure 1.**
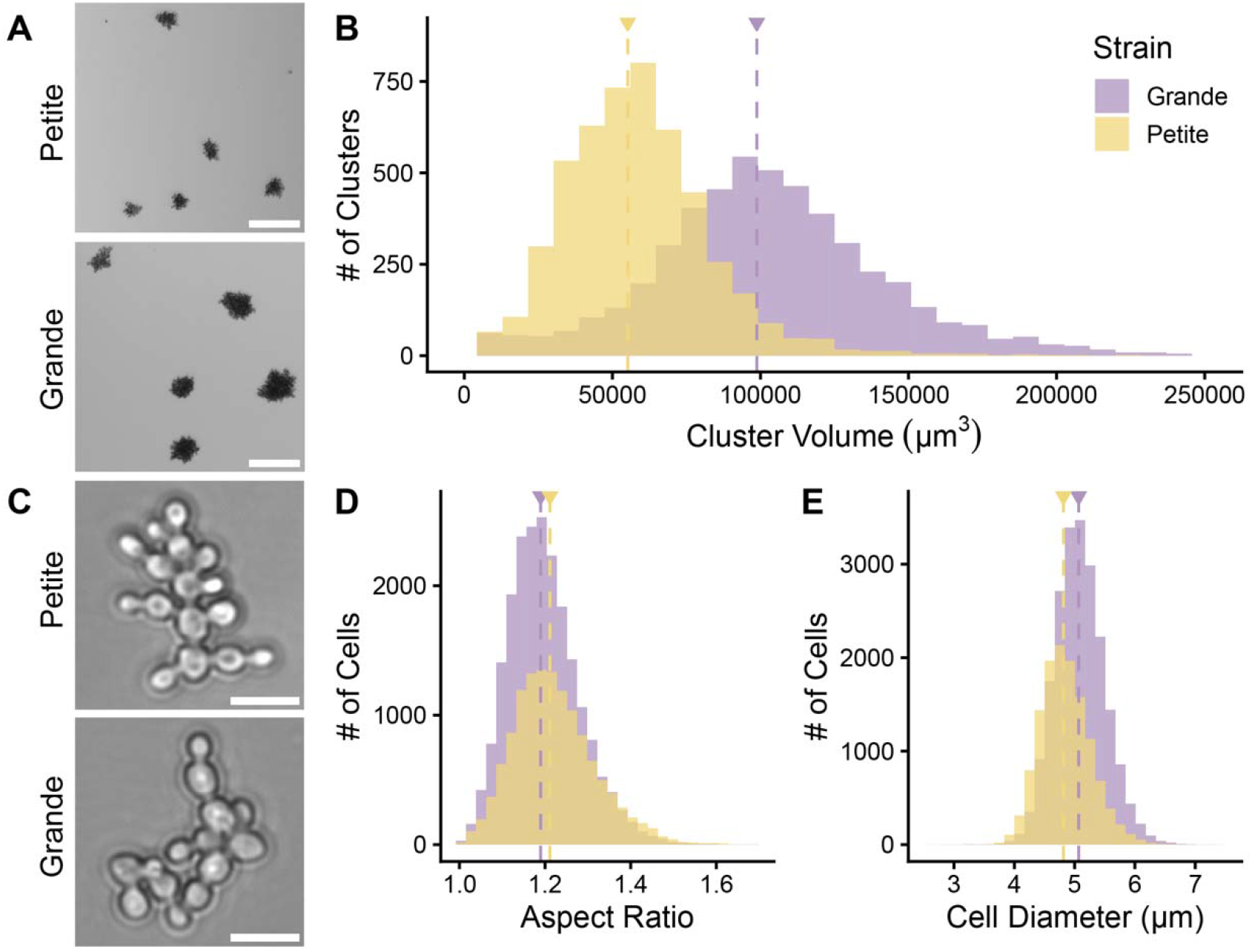
Cell properties cannot explain differences in cluster size between ancestral strains. **A)** Representative images of cluster sizes for the grande and petite ancestors. Scale bar, 100 μm. **B)** Cluster size distributions measured after 2 days of growth for both ancestral strains. Median cluster volumes: grande, 98,756 μm^3^; petite, 55,187 μm^3^ (δ = 0.59, p < 2.2×10^-16^, Wilcoxon test). **C)** Representative images of cells from ancestral strains. Scale bar, 10 μm. **D)** Cell aspect ratio distributions. Median aspect ratios: grande, 1.19; petite, 1.21 (δ = −0.14, p < 2.2×10^-16^, Wilcoxon test). **E)** Cell diameter distributions. Median diameters: grande, 5.07 μm; petite, 4.81 μm (δ = 0.33, p < 2.2×10^-16^, Wilcoxon test). All data were collected from 10 technical replicates grown for 2 days with a 1:100 dilution after the first day.

The grande strain produces significantly larger clusters than the petite strain (**Figure 1A,B**; Cliff’s delta (*δ)* = 0.59, p < 2.2×10^-16^, Wilcoxon test), yet they exhibit relatively minor and opposing differences in cellular aspect ratio (**Figure 1C,D**; δ = −0.14, p < 2.2×10^-16^, Wilcoxon test) and cell size (**Figure 1C,E**; *δ* = 0.33, p < 2.2×10^-16^, Wilcoxon test). Previous studies using snowflake yeast suggested that changes in maximum cluster size are largely explained by these two cell-level phenotypes; increases in either trait are expected to yield increases in maximum cluster size (24,28). This indicates that our current understanding of the determinants of cluster size is incomplete and that additional factors must contribute to the observed size differences. To address this discrepancy, and to develop a more comprehensive understanding of how cell-level properties influence group-level phenotypes in the snowflake yeast system, we devised a soft-agar time-lapse microscopy method to collect data on the following parameters: cellular aspect ratio, cell size, budding angle, and cellular doubling time. To explore the possibility of differences in these phenotypes related to replicative age, we also employed lineage tracing using single-cell tracking to record mother-daughter relationships.

### Single-cell phenotyping by time-lapse microscopy

Initial characterization of the grande and petite strains by time-lapse microscopy (**Figure 2A,B**) showed differences in cellular aspect ratio and cell diameter that were qualitatively similar to our prior static measurements (**Figure 1D,E**; **Figure S1**). Cellular aspect ratio was 1.72% greater in petite cells (**Figure S1A**; Median aspect ratio of petites = 1.18, grande = 1.16; *δ* = 0.01, *p* = 0.89, Wilcoxon test) and cell diameter was 20.12% greater in grande cells (**Figure S1B**; Median cell diameter for grandes = 5.97, petite = 4.97; *δ* = 0.80, *p* = 1.25×10^-11^, Wilcoxon test). We also observed that petite cells bud at 35% wider angles compared to grande cells (Median bud angle = 47.4° in petites vs. 35.2° in grande; *δ* = −0.11, *p* = 0.027, Wilcoxon test. A difference in the angle of attachment between mother and daughter cells could impact the packing density of cells in the cluster, making it a reasonable candidate trait for investigation. Unfortunately, because our observations were limited to only the first 2-3 cell divisions, we cannot conclude whether or not this difference would persist after further cell divisions have occurred.

**Figure 2.**
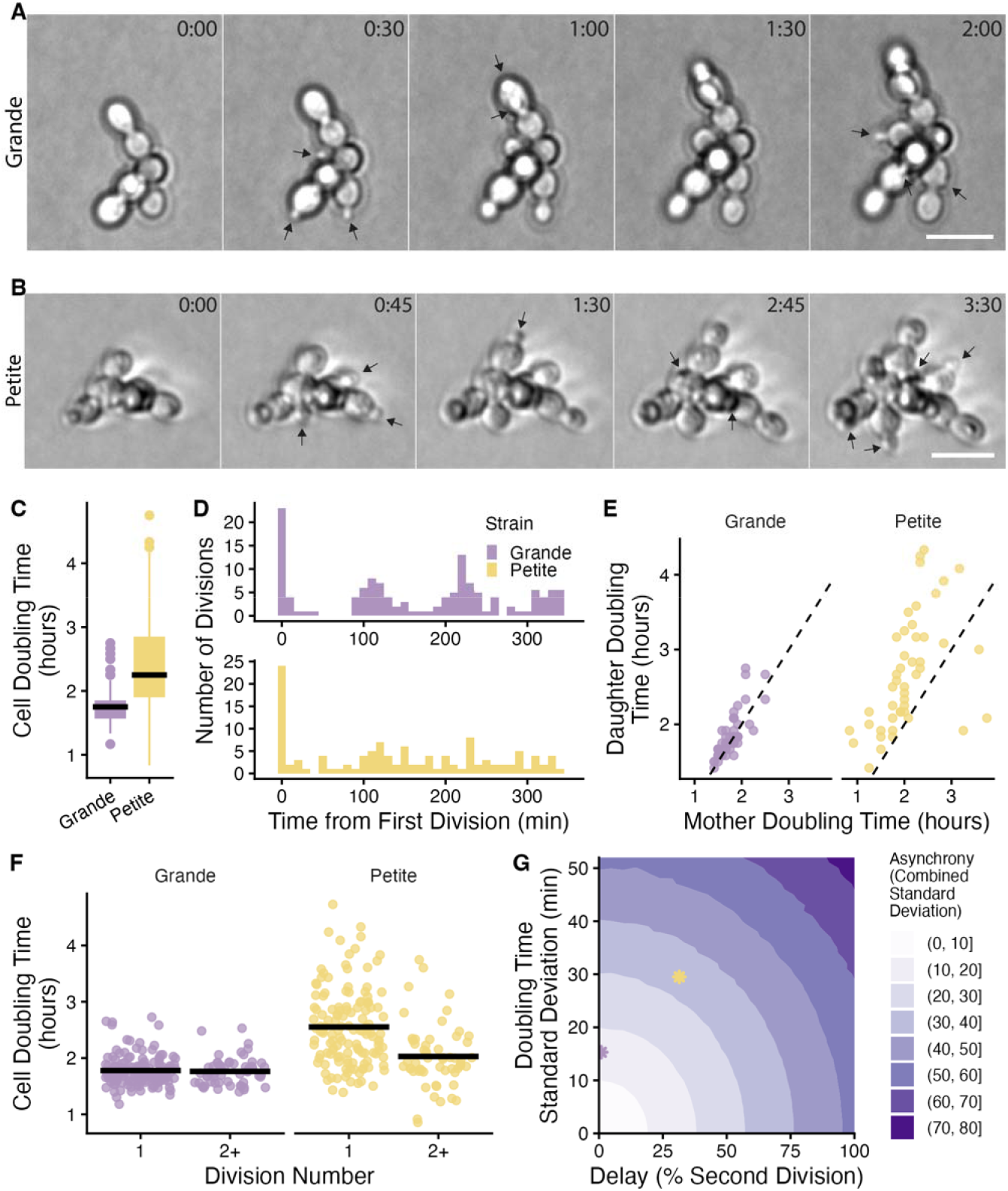
Sources of cell division asynchrony in Ancestral strains of the MuLTEE. **A-B)** Soft-agar time-lapse of the petite and the grande strains (scale bar: 10 µm). **C)** Doubling time distributions of the grande (n=203) and petite (n=191) ancestor strains. Petite strain exhibits greater variation in doubling times compared to the grande ancestor (standard deviation of 41 vs. 15 minutes; p < 2.2×10-16, Fligner-Killeen test). **D)** Histogram of division times with the first division for each cell lineage adjusted to t=0. The “burstiness” of cell division times in the grande strain is suggestive of synchronous cell divisions while the more uniform distribution of cell division times in the petite strain suggests asynchronous cell divisions. Histogram bin size = 10 minutes. **E)** Scatter plots of cell doubling times for mother-daughter pairs for the grande (n = 53) and petite ancestor (n = 46). The grande ancestor divides nearly synchronously; mother and daughter cells have median doubling times only a 5 minutes apart (p = 0.006, r = 0.4; Wilcoxon signed-rank test) while the petite ancestor shows asynchronous division with daughters dividing 35 minutes slower than mothers (p = 2.6×10-6, r = 0.69; Wilcoxon signed-rank test). Dashed diagonal line represent the identity line. **F)** Doubling time distributions of the first and subsequent cell divisions. The petite ancestor exhibits a longer delay in first division (Median of 150 and 117.5 minutes for first and subsequent divisions, respectively; δ = 0.46, p = 3.7×10-7, Wilcoxon test) compared to the grande ancestor (median of 105 minutes for both first and subsequent divisions; δ = 0.02, p = 0.79; Wilcoxon test). Measurements included 147 first and 56 second divisions for the grande ancestor and 135 first and 56 second divisions for the petite ancestor. Horizontal black line represents the mean doubling time. **G)** Heatmap showing how delay and doubling time variation independently contribute to the combined standard deviation of first and second division times (in minutes). A 100% increase in delay produces the same increase in asynchrony as increasing the standard deviation from 0 to 52 minutes. Purple and yellow asterisks represent the delay and standard deviation of the grande and petite doubling time distributions, respectively.

For time-lapse data, cellular doubling time was calculated as the interval between successive bud emergences, which corresponds to the period between consecutive S-phase initiations in yeast (29,30). We found that the petite strain has a 29% longer doubling time than the grande strain (**Figure 2C**; 135 minutes vs. 105 minutes; *δ* = −0.63, *p* < 2.2×10^-16^, Wilcoxon test) and takes longer to bud after the cell has stopped increasing in volume ( **Figure S1B,** suggesting that petites spend longer in G1 phase). Additionally, inspection of the cell division patterns of the petite and grande strains revealed another difference: the cells within clusters of the grande strain divide synchronously while petite cell divisions are more asynchronous (**Figure 2A,B**; **Supplementary Videos 1, 2**; **Table S1**). By plotting the time at which cell divisions occurred relative to the time of the first observed cell division within each cell lineage, we saw periodic bursts of cell division through time for the grande strain, while the timing of cell divisions in the petite strain was more uniform (**Figure 2D**).

Next, we began investigating the source of the asynchronous division pattern we observed in the petite strain. We reasoned that asynchrony could be caused by higher overall variance or by age-structured differences in cellular doubling times. We first determined whether the petite strain simply exhibited greater variation in cellular doubling time than the grande strain. The standard deviation of cellular doubling times for the petite strain was significantly greater than that of the grande strain (**Figure 2C**; 41 vs. 15 minutes; *p* < 2.2×10^-16^, Fligner-Killeen test). To test whether mother cells and their daughters divide at different rates, we measured the similarity of cellular doubling times for mother-daughter pairs of cells for each strain (specifically, we calculated the median difference in doubling time of every daughter cell’s first division and that of their mother’s next division). We found that daughters divided 5 minutes slower than their mothers in the grande strain (**Figure 2E**; *p* = 0.006, *r* = 0.4, Wilcoxon signed-rank test), but petite daughter cells divided 35 minutes slower than their mothers, suggesting a seven-fold increase in the first division delay (**Figure 2E**; *p* = 2.6×10^-6^, *r* = 0.69, Wilcoxon signed-rank test).

To expand our dataset, we compared the distribution of all first divisions (corresponding to the daughter above) vs. all subsequent divisions (corresponding to mother above) regardless of whether we had mother-daughter pairs. All divisions beyond the first are grouped together as “subsequent divisions” because only the first division follows a distinct distribution ( **Figure S2A,B**). With this larger dataset, we observed that the first division is 28% longer than subsequent divisions for the petite strain (**Figure 2F**; 150 vs. 117.5 minutes; *δ* = 0.46, *p* = 3.7×10^-7^, Wilcoxon test), consistent with the difference observed using mother-daughter pairs. In contrast, no difference was observed between first and subsequent divisions for the grande ancestor (**Figure 2F**; 105 vs. 105 minutes; *δ* = 0.02, *p* = 0.79, Wilcoxon test). In the discussion that follows, we will thus refer to the grande strain as “synchronous” and the petite as “asynchronous”.

The pattern observed in the grande strain is consistent with prior work suggesting that yeast lacking *ACE2* failed to exhibit the daughter-specific G_1_ delay that is typically observed for *S. cerevisiae* (31,32). In wild-type cells, Ace2 is thought to play a central role in the regulation of cell size at the Start checkpoint by repressing expression of the G1 cyclin *CLN3*. Deletion of *ACE2* eliminates repression of *CLN3*, resulting in daughter cells that proceed through G1 at the same rate as mother cells. However, both the grande and petite strains of our multicellular yeast lack the *ACE2* gene (as mentioned above, we deleted this gene to engineer the multicellular phenotype because incomplete cell separation is another consequence of *ACE2* deletion) (33,34). Thus, some additional factor must be at play in the petite yeast to reintroduce the daughter-specific delay. For instance, delayed G1-to-S transition has been previously reported in mitochondrial DNA-deficient yeast (35–37). Alternatively, it may be the case that the slower growing petite strain produces smaller daughter cells that need more time to grow before they reach the critical size necessary for Start (38–41). Further work will be required to determine the cause of the first division delay in our system.

To visualize the joint effects of increasing doubling time distribution variation and first division delay on asynchrony, we created synthetic doubling time distributions drawn from log-normal distributions for the first and subsequent divisions with standard deviation of doubling times proportional to the mean of each distribution (see *Log-normal distribution parameterization* in Methods). Log-normal distributions were chosen because they capture the skewed nature of cellular doubling time distributions, with shorter tails for fast divisions and longer tails for slow divisions (42,43). In total, we simulated 1,166 combinations of variation and delay, sampling 10,000 data points per parameter combination. We found that increased asynchrony, defined here as the standard deviation of the overall doubling time distribution, can be caused by either a delay of the first cell division or by increased variance in doubling time (**Figure 2G**). We also confirmed via simulation that either increased variance in cellular doubling times or increased first division delay is sufficient to eliminate the synchronous bursts of cell division over time that we saw with the grande strain (**Figure S3; Figure 2D**). The petite strain exhibits both higher variation and delay than the grande strain (**Figure 2G**, yellow vs. purple asterisk). Thus, in the text that follows we will examine whether these factors, either individually or in combination, could be responsible for the observed disparity in cluster size of the petite and grande strains.

### Cellular behavior alters group morphology in a simple network model

Two distinct features are apparently contributing to the asynchronous pattern of cell division that we observe in the petite yeast: higher variance in doubling time and a delayed first cell division. To determine whether either, or both, of these factors could be contributing to the unexplained discrepancy in cluster size that we observed between the petite and the grande strains, we developed a simple network model of snowflake yeast, where cells are represented as nodes and mother-daughter connections as edges. Each cluster begins as a single cell and all daughters remain attached to their mothers after division. To enable first division delay and doubling time variation, cells reproduce according to user-defined “first division” and “subsequent division” doubling time distributions that can exhibit differences in mean and standard deviation. To isolate the effects of cell division timing on group structure, we ignore the potential impacts of nutrient diffusion (26,44,45) or growth limitations that may arise due to crowding (46,47).

With this simple model of cluster growth, we can study a number of topological properties relevant to fitness in the snowflake yeast system. First, the diameter of a cluster can be approximated using the “network diameter”, defined as the length of the longest shortest path between all possible pairs of nodes in the network. Furthermore, we can obtain a proxy for the local packing density of cells (and the associated physical strain on any given cell-cell connection) by calculating the “edge degree” for each edge in the network as the sum of other edges emanating from the two connected nodes, excluding their mutual connection. As we will detail in the next section, the accumulation of physical strain plays a critical role in determining the timing and location of fragmentation in snowflake yeast (28).

Using three distinct doubling time distributions, we simulated network growth from a single cell to 1300 nodes, which represents the maximum number of cells measured in a grande ancestor cluster (**Figure S4A,B**). These distributions included: the empirical petite ancestor data (high variation, first division delay), the empirical grande ancestor (low variation, no delay), and a synthetic distribution matching the petite ancestor but without delay (high variation, no delay) (**Figure 3A**). Together, these three distributions enable us to isolate the effects of delay from the effects of variation. At a network size of 100, network diameter was ∼8.3% smaller only for networks generated using the petite strain’s doubling time distribution (**Figure 3B**; p < 2.2×10^-16^ for petite vs others, and p = 0.07 for petite w/o delay vs grande, Wilcoxon test). Similarly, we observe that maximum edge degree in 100-node networks generated using the petite strain’s doubling time distribution was ∼9% greater (**Figure 3C**; p < 2.2×10^-16^ for petite vs others, and p = 0.24 for petite w/o delay vs grande, Wilcoxon test). This suggests that the delayed first division can lead to more highly branched networks with shorter diameters. Higher variance in doubling time alone (petite without delay distribution), however, does not produce a shift in the mean of either phenotype.

**Figure 3.**
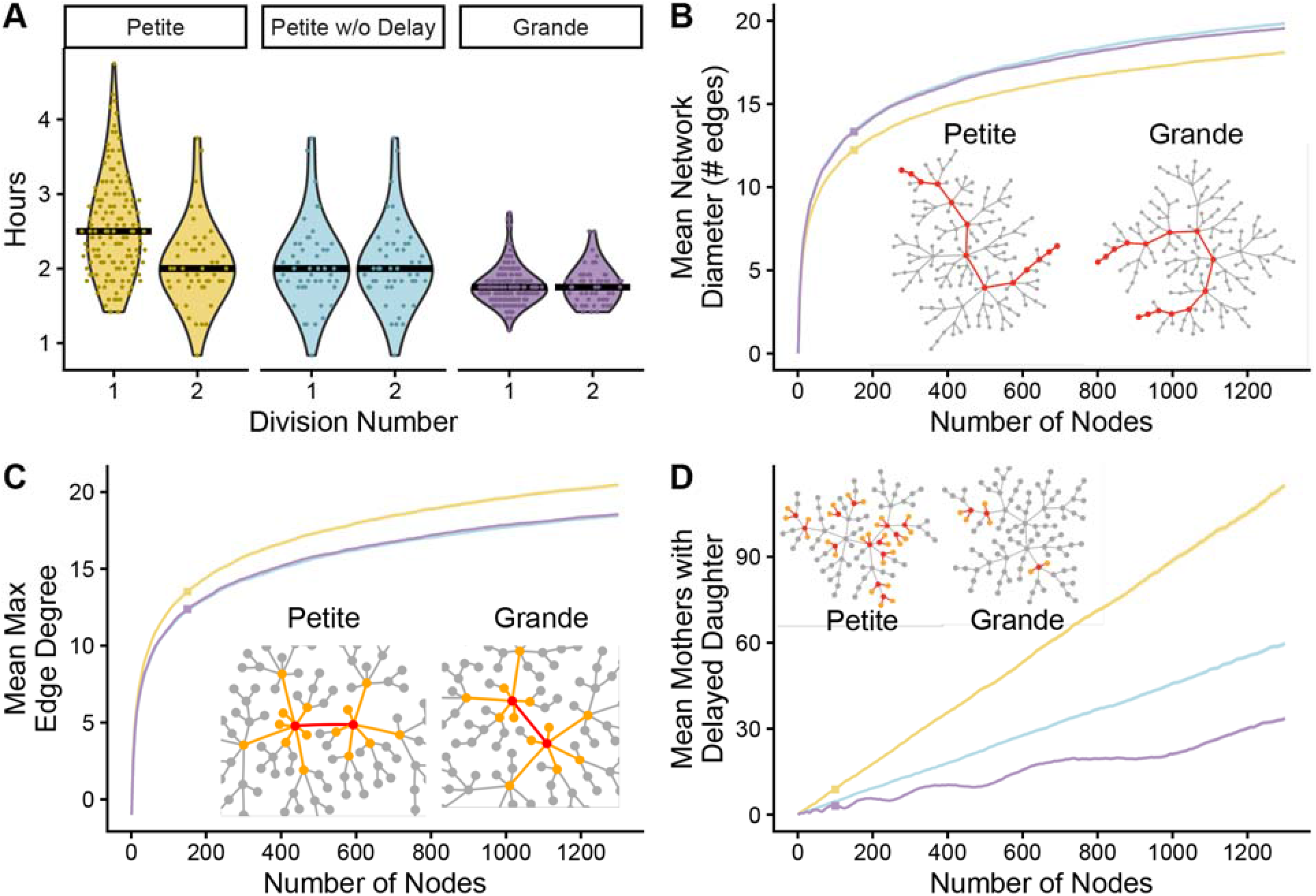
Cell behavior affects group morphology by changing network topology. **A)** Doubling time distributions used in simulations: petite ancestor (high variation with first division delay), modified petite ancestor (high variation, no delay), and grande ancestor (low variation, no delay). **B)** Mean network diameter by network size. Inset: Diameter-defining paths (red nodes and edges). **C)** Mean maximum edge degree by network size. Edge degree is defined as the sum of node degrees for both nodes forming an edge minus 2 (excluding their mutual connection). Inset: Maximum edge degree connections (edge of interest in red, contributing edges in orange). **D)** Mean number of mothers with ≥2 unbudded daughter cells as a function of network size. Inset: Motifs of mother cells with ≥2 unbudded daughters (mothers in red, daughters in orange). For panels **B-D**: Each trajectory shows the average of 300 independent simulations with network properties measured after every 5 nodes added. Square markers indicate network sizes shown in insets. Example networks shown have diameters of 12 and 13 (petite vs. grande, panel **B**), edge degrees of 14 and 12 (panel **C**), and 13 and 3 motifs (panel **D**), respectively.

By comparing 100-node network topologies, we also observed that mother cells with two or more unbudded daughter cells were 2.8 and 1.7 times more common in petite networks compared to the grande strain and the petite without delay strain, respectively (**Figure 3D**, red nodes connected to orange; δ = 0.93 vs. grande, δ = 0.78 vs. petite w/o delay, p < 2.2×10^-16^ for both comparisons, Wilcoxon test). These “unbudded daughter” motifs, which arise when a mother divides faster than its daughter, are of interest because they typify the phenotypic consequences of asynchrony: they produce an increase in edge degree without increasing the overall network diameter. Grande networks also produced 40% fewer unbudded daughter motifs than the petite without delay condition (**Figure 3D**; δ = 0.46, p < 2.2×10^-16^, Wilcoxon test), suggesting that both first division delay and higher variation in doubling time contribute to the appearance of these motifs. However, because the asynchrony of the petite without delay condition is essentially unbiased, these unbudded daughter motifs are offset by an equal number of “filamentous” motifs where daughters divide faster than their mothers. High variance in doubling times can therefore have profound effects on the topology of networks while having little to no effect on some group-level phenotypes (explored further in **Figure 8**).

### Linking morphological changes to multicellular life cycle consequences

To understand how these morphological differences translate into consequences for the snowflake yeast life cycle, we extended our network model to capture both the growth and reproduction of clusters. In snowflake yeast, group reproduction occurs when a single cell-cell connection is broken due to crowding-induced mechanical stress (28), hence a dynamic model of fragmentation requires the specification of both when and where fragmentation occurs. Biophysical modeling has been successful in predicting the timing of fragmentation, but not the spatial location (23,48). While solving this prediction problem in an explicit biophysical model is outside of the scope of this work, we reasoned that we could use our edge degree metric to approximate the local accumulation of crowding-induced mechanical stress on cell-cell connections. Specifically, we assume that each edge in the network has a maximum edge degree threshold at which it can no longer maintain the cell-cell connection (this approach is conceptually similar to the “kissing number” parameter in (Nanda et al., 2024)). Fragmentation therefore occurs when an edge has a degree that meets or exceeds this threshold (timing) and the edge that is severed is the one that crossed the degree threshold (location). These simple assumptions localize fracture events to the most densely connected regions of the network and the size at fracture can be tuned by adjusting the edge degree threshold. To select the edge degree threshold for life cycle simulations, we calibrated our model using the grande ancestor’s fracture size (Maximum edge degree = 15, Fracture size = 338; **Figure S4C**). Using this dynamic network modeling framework, and this experimentally calibrated edge degree threshold, we simulated population growth across eight cluster generations for each strain type shown in **Figure 3A**, tracking key properties at each fragmentation event.

First, we measured the average cluster size at fracture. Both synchronized strains achieved substantially larger fracture sizes than the petite ancestor (62% larger for petite without delay and 55% larger for grande; Figure 4A), with the petite without delay reaching a larger size than the grande ancestor (354 vs. 339.4 nodes; further explored in **Figure 5**). We then compared our results to an existing biophysical model where cells are represented as ellipsoids of defined diameter and aspect ratio and mechanical stress relate to the cumulative overlap of cells in space (23,24,48). To incorporate the differences in cell division timing, we adapted the model to add cells in the same temporal sequence as our network simulations, with fragmentation checks occurring only after all cells from each time point were added (**Figure S5A-C**). We again found that clusters generated using the doubling time distribution of the petite contain 7.8% fewer cells and have 19% less volume than those generated using the grande strain’s doubling time distribution (**Figure S5D,** 312.7 vs. 339.3 cells; **Figure S5E,** 28,976 vs. 35,983 µm^3^). The effect of doubling time delay is qualitatively similar to the effect observed in the dynamic network model, but smaller in magnitude (27 cells difference in the biophysical model compared to 121 cells in the network model). Furthermore, it provided independent support of the idea that asynchronous division accelerates mechanical stress accumulation through more localized crowding around older cells (**Figure S5F**, distance between overlaps = 13.26 µm vs. 13.92 µm).

**Figure 4.**
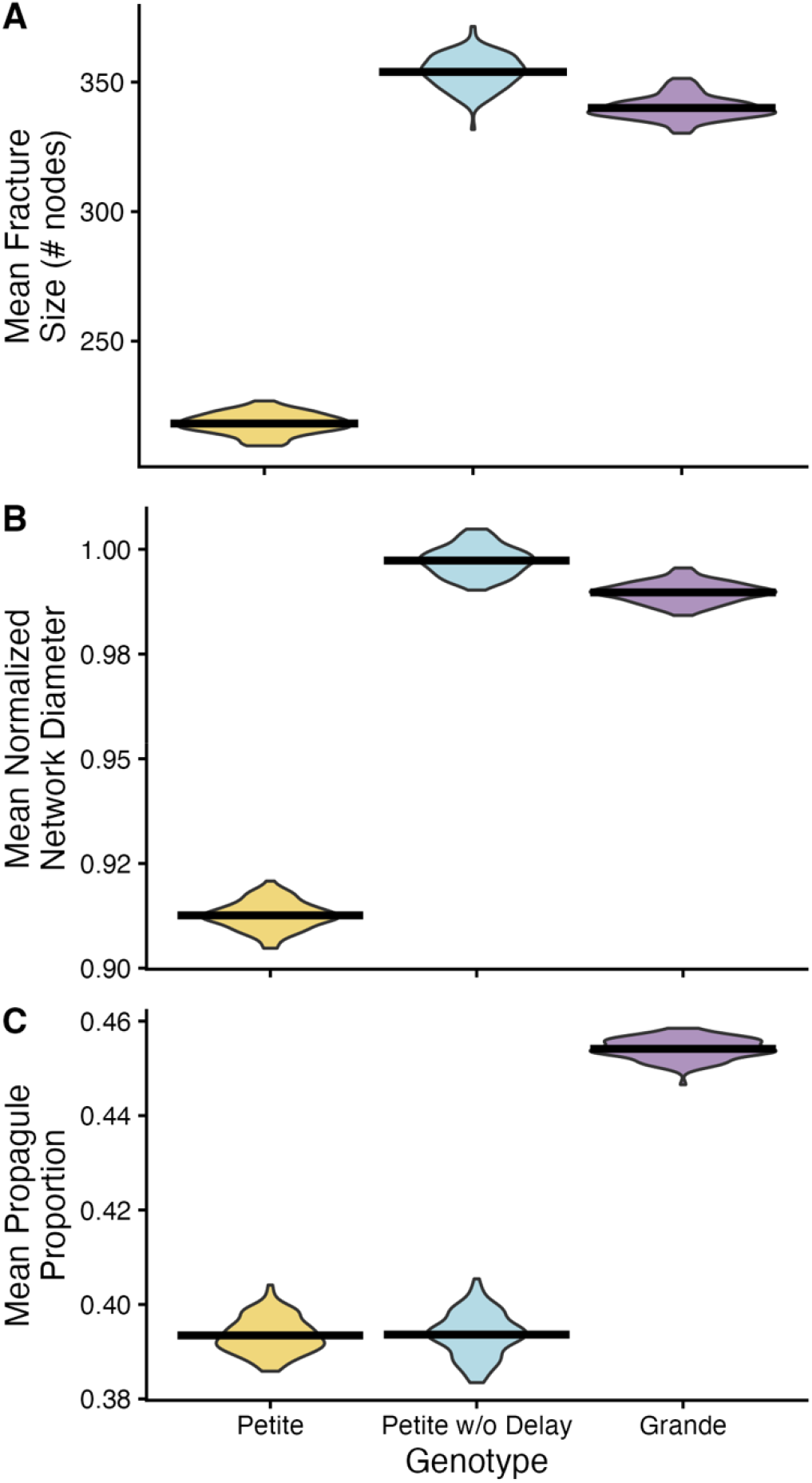
Delayed first division results in smaller, more branched multicellular groups at fracture and asynchronous cell division results in smaller propagules. **A)** Cluster fracture size distributions for the three strains. Median sizes: petite = 218.3, petite without delay = 354.0, grande = 339.4. All means significantly differ (pairwise Wilcoxon tests with Bonferroni correction, p < 2e-16). **B)** Normalized network diameter relative to an exponentially growing network. Values >1 indicate higher diameter than in an exponential network, <1 indicate lower diameter. Medians: petite = 0.912, petite without delay = 0.997, grande = 0.990. All medians significantly differ (p < 2e-16, Wilcoxon test with Bonferroni correction). **C)** Propagule proportion after fracture (propagule nodes divided by nodes of the cluster before fracture). Medians: petite = 0.393, petite without delay = 0.393, grande = 0.453. Grande ancestor differs from both of the other strains (*p* < 2.2×10^-16^, for grande vs petite and grande vs petite w/o delay, and petite vs petite w/o delay *p* = 1, Wilcoxon test with Bonferroni correction). For all these results 100 simulations were performed by allowing the clusters to grow up to 8 fragmentation events (8 cluster generations), and the plots show the mean value obtained from all the clusters in one simulation.

**Figure 5.**
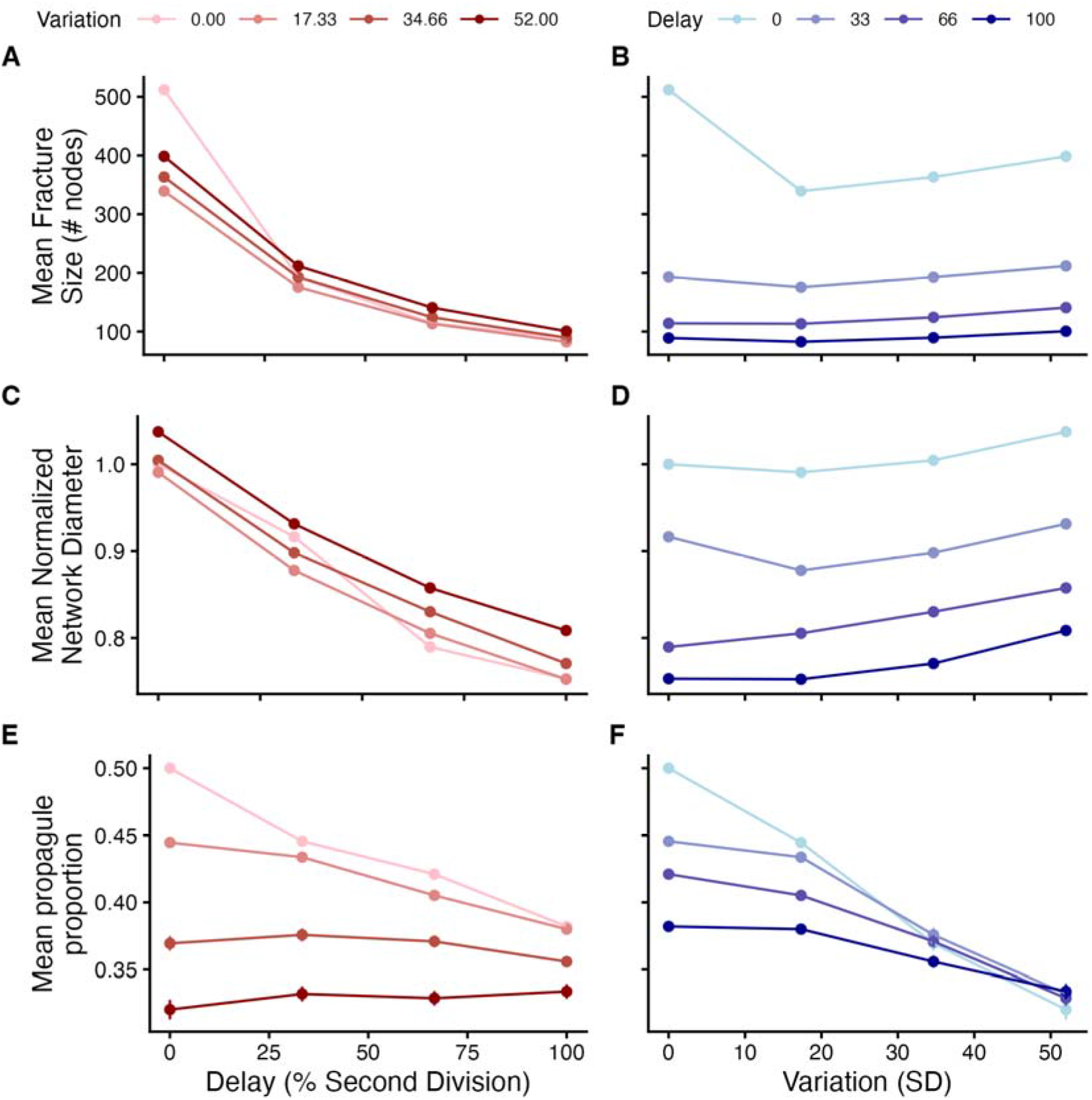
Effects ofdelay and variation in doubling time distributions on cluster properties. Effects of 4 different levels of delay and variation on mean cluster fracture size **(A, B),** mean network normalized diameter **(C, D)**, and mean propagule proportion **(E, F)**. Left column highlights changes in first division delay and right column highlights changes in variation of the doubling time distribution (represented by standard deviation in minutes). Each dot is the grand mean of 30 simulations. Each simulation consisted of 50 sequential cluster fragmentation events where fragmentation is followed by growth of either the propagule or parent at random.

The difference in cluster size at fracture is driven by differences in network topology. Cells with multiple undivided daughters contribute disproportionately to reaching the edge degree threshold without increasing overall group size. To analyze the effects of network topology on fragmentation, we normalized network diameter to that of an exponentially growing network, enabling comparison across different cluster sizes (Figure S6). This normalized metric reveals underlying structural patterns: values below 1 indicate networks enriched by undivided nodes (with degree 1), while values above 1 reflect configurations with fewer undivided nodes and more degree-2 nodes. The petite ancestor exhibited a normalized diameter of 0.912 (Figure 4B), consistent with its first division delay pattern. In these clusters, the delayed initial division followed by rapid subsequent divisions creates highly connected older cells. In contrast, strains without first division delay showed diameters closer to 1, indicating network topologies more closely resembling exponential growth patterns. Normalized network diameter was ∼8.6% and ∼9.3% greater for the grande ancestor (0.990) and petite without delay strain (0.997), respectively (Figure 4B; p < 2e-16, Wilcoxon test with Bonferroni correction). All properties shown in Figure 4 remained constant through time in a modified simulation that tracked a single randomly-chosen cluster after fragmentation over 100 cluster generations (Figure S7), further indicating that changes to the doubling time distribution result in characteristic phenotypic consequences.

We also predicted that changes to the timing of cell divisions could affect the size of propagules produced by fragmentation. Propagule size has important direct and indirect fitness effects for nascent multicellular organisms (46,49–51). We define the propagule as the smaller of the two clusters resulting from a fragmentation event and “propagule proportion” as the ratio of cells in the propagule to the total number of cells in the pre-fragmentation cluster. Accordingly, this metric ranges from 0.5, for symmetric division, to near 0 when propagules contain just a few cells and, in our model, only an exponentially dividing network will achieve perfect symmetric division. Notably, the petite and petite without delay clusters have propagule proportions of 39.3% while the grande ancestor achieves more symmetrical fragmentations with propagule proportions of 45.3% (**Figure 4C**, *p* < 2.2×10-16, for grande vs petite and grande vs petite w/o delay, Wilcoxon test), suggesting that high variation in doubling time distribution alone is sufficient to explain the decrease in propagule proportion for the petite strain. This differs from our earlier findings on network diameter and maximum edge degree where high variance in cellular doubling time did not contribute to shifts in the mean of cluster-level phenotypes. Propagule proportion is uniquely affected by increases in variation because any deviation from perfect exponential growth will disrupt the symmetry of fragmentation.

To systematically dissect how division timing shapes network topology, we simulated cluster growth using synthetic doubling time distributions drawn from log-normal functions. We varied first division delay and temporal variation, sampling from a range of four distinct values for each parameter which had comparable effects on asynchrony (Figure 2G). Our analysis revealed that lower first division delay was associated with higher size at fracture and higher mean normalized diameter, while differences in doubling time variation had no significant effect on these properties (Figure 5A-D). The lower size at fracture of the grande ancestor compared to the petite without delay (Figure 4A) can be attributed to the small negative effect of decreasing variation when there is no first division delay (Figure 5B; compare variation = 17.33 to variation = 34.66 when delay = 0, light blue trajectory). Notably, propagule proportion responded to both timing parameters, with symmetric fragmentation emerging only when both variation and delay were minimal (Figure 5E, F). This explains why only the grande ancestor achieves near-symmetric division with propagule proportions approaching 0.5 ( Figure 4C), as eliminating delay alone is insufficient to ensure symmetrical fragmentation.

### Partitioning the selective effects of first division delay

Having established that first division delay significantly affects the size of clusters at fragmentation, we next investigated the impact this change could have on relative fitness under experimental conditions similar to those used in the MuLTEE. Each passage of the MuLTEE involves a 24-hour growth phase in liquid media followed by a brief phase of “settling selection” in which a subsample of the population is transferred to an Eppendorf tube where the culture is then vortexed and left to settle for 3 minutes before the top ∼95% of the volume is discarded (24). Since the lack of a first division delay is associated with both faster individual growth and larger cluster size at fragmentation (compare petite vs. petite without delay, **Figure 3A**), selection during either the growth phase (where fitness is largely determined by differences in cellular growth rate) or during settling selection (where fitness is largely determined by differences in cluster size) should favor the loss of first division delay (52). To quantify these effects and determine their relative contributions to fitness we simulated competition experiments using our dynamic network model with periodic settling selection based on simulated volumes from our biophysical model.

To investigate the fitness effects of first division delay, we competed networks generated using the doubling time distribution of the petite ancestor strain vs. that of the petite without delay strain. Prior to competition, populations of each strain were grown independently until they reached stable cluster size distributions. Competition assays were then initiated by randomly sampling clusters from these stable distributions until ≥100,000 cells were collected, maintaining an approximate 1:1 ratio of strains on a per cell basis. These cocultures were then passaged for 20 transfers or until we observed the extinction of one of the two strains. Each passage consisted of growth to a carrying capacity of 10 million cells (this was shown to be a sufficiently large population to accurately measure fitness; **Figure S8**) followed by two sequential 10-fold dilutions: the first to mimic the random subsampling of the total culture volume and the second to mimic the passaging of biomass that reached the bottom 5% of the tube after 3 minutes of settling selection (24). Sedimentation rates were calculated using Stokes’ law for all clusters surviving the first 10-fold dilution (24,27,53). Fitness during the growth and settling phases were estimated separately by measuring selection rates as the difference in Malthusian parameters of the two competing strains (54,55). Selection rates from the growth phase of the first transfer were excluded from this analysis, as the initial lower concentration of the fast-growing strain allowed for additional generations of growth compared to subsequent transfers (**Figure S9**).

Under conditions that closely mimicked those of the MuLTEE, we found that while the petite without delay strain was advantaged during both the growth and settling phases of the experiment, the selection rate during growth (*r_G_*) was ∼36% higher than it was during the settling phase (*r_S_*) (**Figure 6A**; *r_G, Standard_*= 0.641, *r_S, Standard_ =* 0.470; *δ* = 0.74, *p* = 1.07×10^-10^, Wilcoxon test). If we eliminate the growth advantage by setting the overall cellular growth rates of the strains with and without delay to be equal, we observe the same fitness benefit during settling selection (**Figure 6B**; *r_S, Standard_* = 0.470, *r_S, Equal_* = 0.496; *δ* = 0.154, *p* = 0.185, Wilcoxon test). This suggests that the benefits of synchronous division during settling selection are conferred by the changes in group morphology and size, not due to a cross-level byproduct of increased cellular growth rate. We then manipulated the strength of settling selection by simulating a modification of the standard settling selection protocol: initiating clusters at the top of the tube rather than at random positions in the media column. Under these altered conditions, the selection rate during settling was ∼533% higher than the selection rate during growth (**Figure 6C**; *r_G, Standard_* = 0.640, *r_S, Standard_* = 4.05; *δ* = −1, p < 2.2×10^-16^, Wilcoxon test), revealing how environmental context can dramatically alter the relative importance of group-level traits in evolutionary outcomes. Again, the fitness benefit during settling selection was not dependent upon differences in cellular growth rate (Figure 6D; r_S, Standard_ = 4.05, r_S, Equal_ = 4.14; *δ* = −0.002, *p* = 0.986, Wilcoxon test).

**Figure 6.**
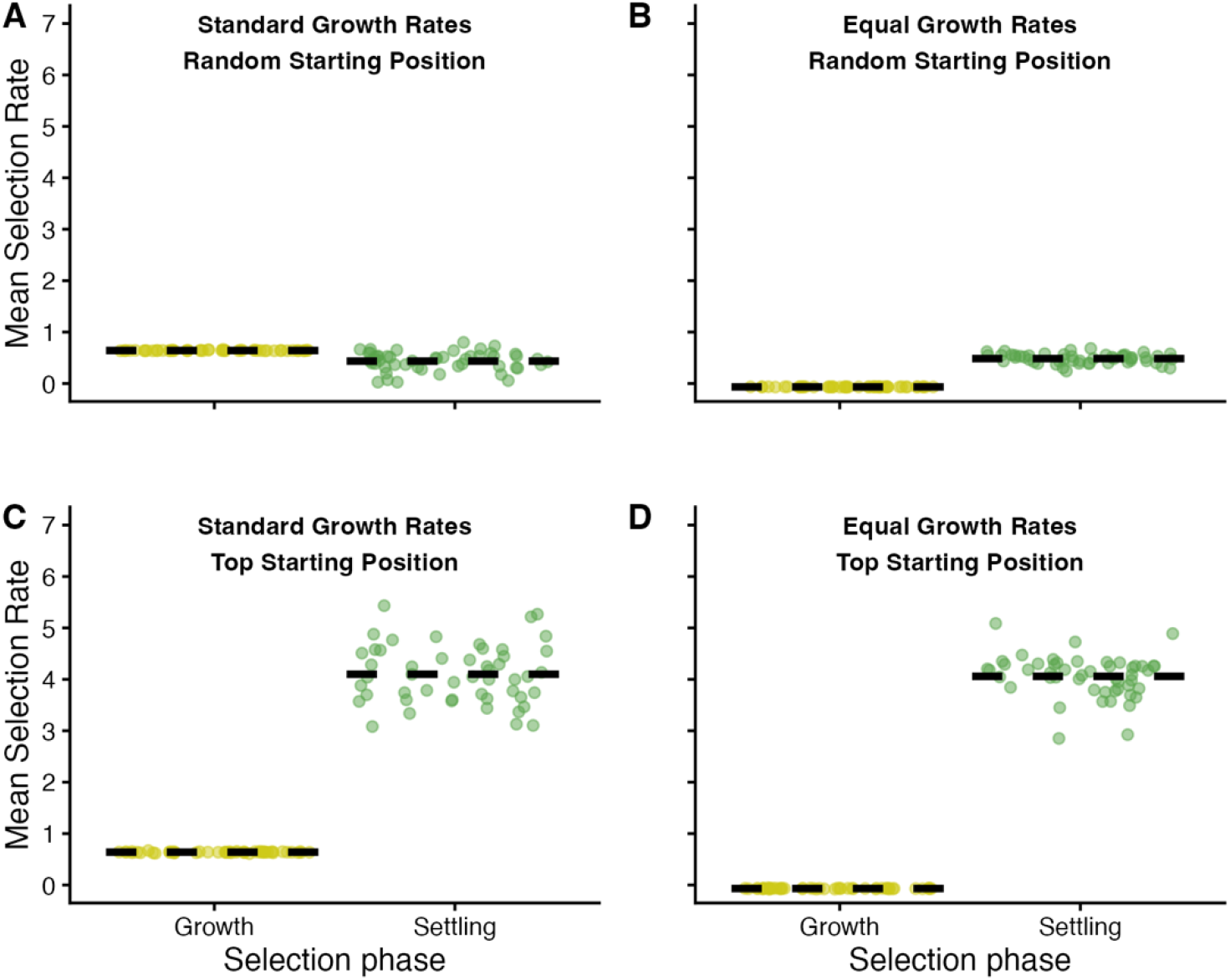
Simulated selection rates for growth and settling phases under different selection regimes and growth rate conditions. Selection rates were calculated from competition simulations using the doubling time distributions of the petite ancestor and the petite without delay strain to assess how environmental conditions and cellular growth rates influence selection during the growth an settling phases of the experiment. Two environmental conditions were tested: clusters starting at random positions (matching experimental conditions, **A** and **B**) or at the top of the tube to maximize settling selection effects (**C** and **D**). Two growth rate scenarios were examined: standard growth rates (**A** and **C**), where the petite without delay has faster overall growth rate, and equal growth rates (**B** and **D**), where the net growth rate of the petite without delay was made equal to the petite ancestor by adding 15 minutes to the average doubling time. Each data point represents the mean selection rate per phase (averaged across up to 20 transfers) for each of the 50 replicate competition experiments.

### Changes in cellular doubling time distributions over 5,000 generations of evolution

Our simulated competitions clearly predict that any mutation that eliminates the first division delay observed in the petite ancestor strain should be selectively favored under the experimental conditions of the MuLTEE. To test this prediction, we performed time-lapse microscopy on the descendants of the petite ancestor after 200, 400, 600, and 1000 days of evolution (corresponding to ∼5,000 generations). Consistent with our predictions, when we compared the first and subsequent doubling time distributions of our evolved strains, we found that the first division delay had been lost in all 5 replicate populations by day 200 (∼1,000 generations) and this remained true through day 1000 (Figure 7; p > 0.05 after Bonferroni correction for all evolved strains; Wilcoxon test). As was the case for our ancestral strains, when we restrict our analyses to mother-daughter pairs, we detect some small but significant differences for 6 of the evolved strains after day 200 (PA1 t400, PA2 t400, PA2 t600, PA3 t600, PA5 t600, and PA1 t1000 showed slight delays in daughter cell division; Figure S10; p < 0.02 after Bonferroni correction, Wilcoxon rank-sum test). However, the differences in mother vs. daughter doubling time were significantly reduced relative to the petite ancestor (p < 0.05 after Bonferroni correction for all evolved strains; one-sided Wilcoxon test); all detected delays were less than 2 minutes on average by t600, a 94% reduction in delay relative to the petite ancestor. While it is tempting to interpret this result as evidence of selection favoring synchronous cell division, it is also possible that synchronous growth is a mere side effect of fast growth and that it appears as consequence of mutations that compensate for the growth defects imposed by the petite mutation. Regardless of the ultimate cause of this change in the MuLTEE, our model shows how manipulation of mother-daughter doubling times can provide a lever for the control of multicellular morphology that can affect fitness.

**Figure 7.**
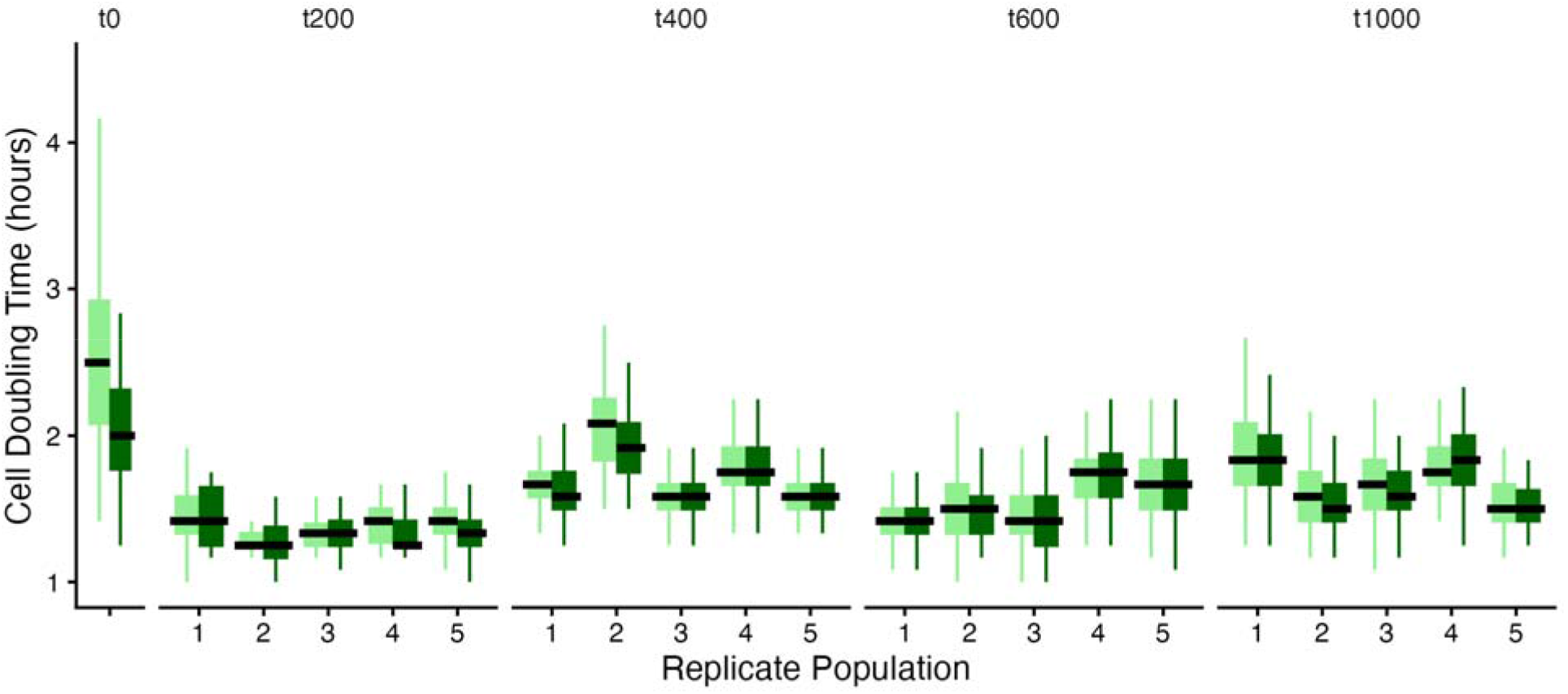
Recovery of synchronous cell division in the evolved derivatives of the petite strain. Doubling time distributions of the first cell division (light green) and subsequent cell divisions (dark green). The petite ancestor strain (t0) is the only one in which the first division is significantly slower than the subsequent division (p = 7.92×10^-6^ after Bonferroni correction, Wilcoxon test). All evolved strains have no statistically significant delay in the first division (p > 0.06 for all evolved strains after Bonferroni correction, Wilcoxon test). **Tables S1** and **S2** report the total number of time-lapses collected and cell doublings times measured.

### Further increases in group size are enabled by accelerated first division

Up until this point, we limited our focus to asynchronous cell divisions caused by a delay in the first cell division because this was the pattern observed in our multicellular yeast ancestors. However, any difference in the mean doubling times for the first and subsequent cell division will lead to asynchrony, including an acceleration in the first division time. Here, we relax this limitation to consider these alternative scenarios. We hypothesized that faster first divisions would result in larger fragmentation sizes, as clusters would expand outward more rapidly, forming more filament-like branches, allowing the cluster to accumulate more cells before reaching the edge degree fragmentation threshold. To test this hypothesis, we simulated cluster growth using synthetic log-normal doubling time distributions with varying levels of first cell division delay, ranging from positive values (as observed in the evolution of multicellular yeast) to negative values representing accelerated first divisions (**Figure 8A**).

**Figure 8.**
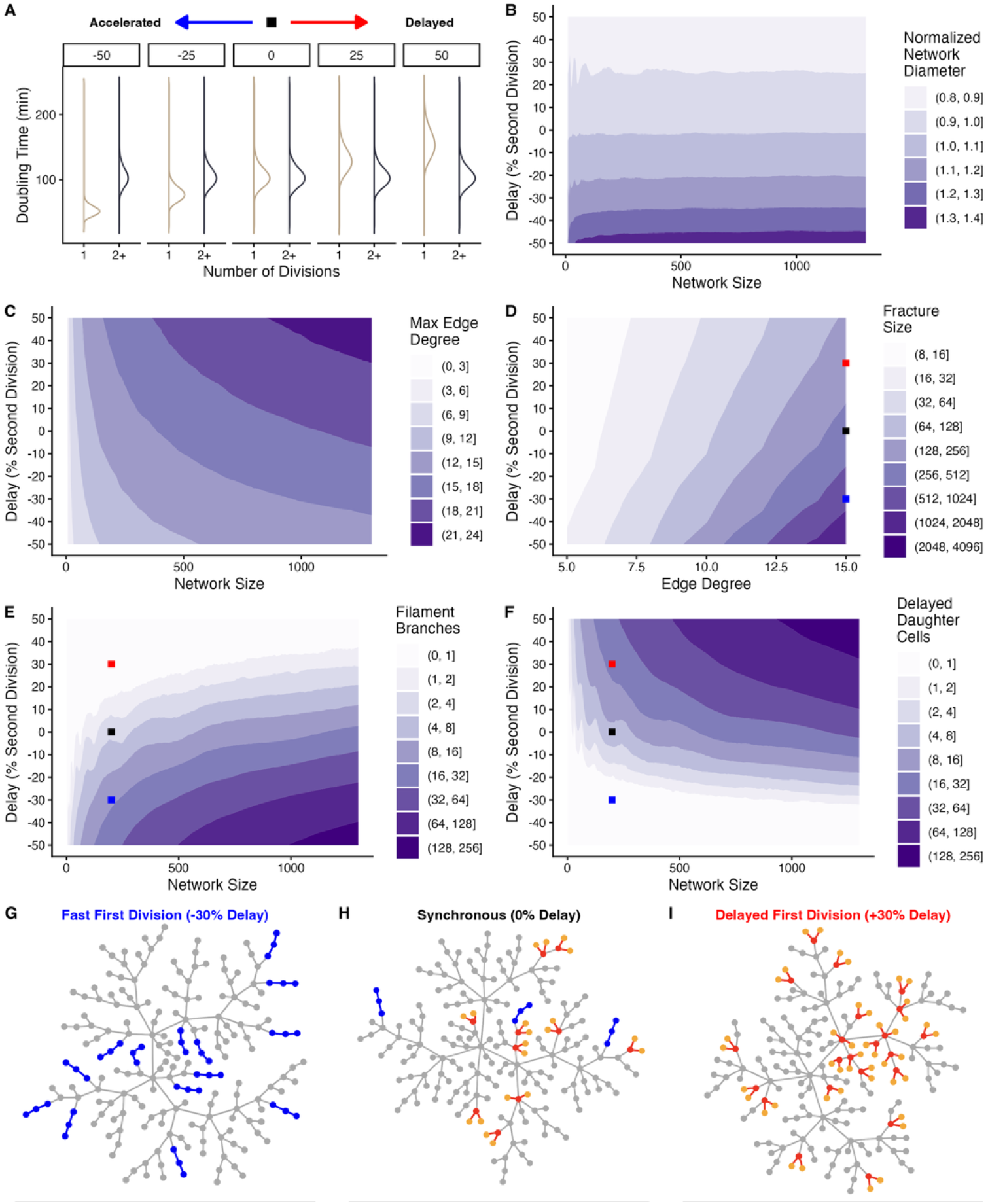
Accelerated first cell division creates a different network topology that allows for the formation of larger clusters. **A)** Synthetic doubling time distributions for strains with fast first division (−50% and −25% delay), synchronous cell divisions (0% delay), and delayed first division (25% and 50% delay). Standard deviation = 15 minutes for all distributions. Simulations tested 21 delay values from −50% to +50% in 5% increments. **B)** Contour plot of mean normalized network diameter. **C)** Contour plot of mean maximum edge degree. **D)** Mean fracture size across different edge degree thresholds. **E)** Contour plot of mean filament branch motifs. A filament branch is defined as at least 3 contiguous connected cells with a node degree of 2 or less. **F)** Contour plot of mean mother cells with ≥1 delayed daughter cell. Color scales are logarithmic in panels **D, E,** and **F**. For panels **B, C, E,** and **F**: 300 simulations per delay value, grown from 1 to 1300 cells. For panel **D**: 30 simulations per edge degree (5-15) and delay combination, tracking fracture size over 50 generations with random parent/propagule selection after fragmentation. Blue, black, and red dots in D-F mark example networks with fast first division (panel **G**), synchronous division (panel **H)**, and delayed first division (panel **I**), respectively. **G)** Example network (200 cells) with fast first division (−30% delay). **H)** Example network (200 cells) with no delay (synchronous). I) Example network (200 cells) with delayed first division (30% delay). Colored nodes represent motifs described in panels **G − I**; blue nodes represent filamentous branches where daughter cells divided faster than their mothers, red (mothers) and orange (daughters) highlight motifs of mother cells with ≥2 undivided daughters.

Extending our previous results, we find that delay is negatively correlated with normalized network diameter (**Figure 8B**; the normalization function makes this effect independent of network size) and positively correlated with maximum edge degree (**Figure 8C**). Accordingly, size at fracture increases exponentially as delay decreases and the impact of this delay compounds as the edge degree threshold increases (**Figure 8D**). For instance, at an edge degree threshold of 15, the mean size at fracture for a network generated with a delayed first division is 184 nodes (**Figure 8D**; red point), while synchronous and accelerated first division distributions produce a mean size at fracture of 337 and 829 nodes, respectively (**Figure 8D**; black and blue points). Finally, as hypothesized, networks generated with faster first divisions exhibited a greater number of “filamentous” network motifs, where daughter cells divide faster than their mothers (**Figure 8E**) while the previously characterized “unbudded daughter” motif was more prevalent in slow first division networks (**Figure 8F**). While networks with severe differences in first and subsequent division times may exhibit only one type of these motifs (e.g., exclusively filamentous motifs when delay is strongly negative and viceversa), both motifs will often occur in a single network when variance in doubling time is sufficiently high relative to delay (**Figure 8G-I**). This combination of both types of motifs causes the mean of the normalized network diameter of synchronous networks to remain close to 1, similar to that of exponentially dividing networks, since the opposing effects of both motifs counterbalance each other.

We next asked whether the striking increase in size at fracture with accelerated first division held up in a biophysical model that takes into account the spatial constraints of three-dimensional growth. Comparing the biophysical and network models, we found no significant difference in the predicted size at fracture for the delayed first division conditions ( **Figure S11**; delay of 25 and 50% of second division; *p* > 0.05 after Bonferroni correction, Wilcoxon test). In contrast, with synchronous division or an accelerated first division, the biophysical model predicted sizes at fracture that were 6%, 46%, and 79% smaller than the network model for delays of 0%, −25%, and −50%, respectively (**Figure S11**; delay = 0%: 311 vs. 332 cells; delay = − 25%: 374 vs. 692;5 cells; delay = −50%: 446 vs. 2120.5 cells; *p* < 0.05 after Bonferroni correction, Wilcoxon test). This suggests that our simple edge degree threshold criterion for fragmentation works best while clusters are relatively small (*i.e.*, when cell division is nearly synchronous or delayed), but breaks down as clusters increase in size and cellular packing poses a major problem. However, while the network model overestimates the size of the effect, both models predict that an accelerated first division will result in an increased size at fracture (**Figure S11**; *p* < 2×10⁻¹^□^, Wilcoxon test). Overall, the network model with the maximum edge degree threshold for fragmentation reproduces the biologically observed cases and the qualitative trend of the more complex biophysical model.

### Apoptosis provides an alternative path to large group size

Given the predicted increase in cluster size via accelerated first division in our network and biophysical models (**Figure 8; Figure S11**), it is somewhat surprising that we do not observe accelerated first division evolving in the MuLTEE (**Figure 7; Figure S10**). However, we do observe the repeated, parallel evolution of another cell-level trait that could have similar impacts on cluster morphology, albeit via an entirely different mechanism: elevated rates of apoptosis (22,56). Elevated rates of apoptosis are thought to have been favored in snowflake yeast because dead cells serve as weak points, facilitating fragmentation and releasing cells from the interior of large clusters where spatial constraints or nutrient diffusion may limit their growth (22,46,57). Another side-effect of cell death is that after a cell dies, it leaves no further offspring. This otherwise mundane and obvious consequence of cell death, taken in the context of our network model of snowflake yeast growth, implies that, for a network of any given size, network diameter will increase as a function of the apoptosis rate. Furthermore, because fragmentation is driven by local packing density of cells, cell death could even increase the size at which fragmentation occurs by creating pockets of less densely packed cells (that is, higher apoptosis rates may lead to a reduction in the average edge degree).

We examined the effects of apoptosis, independent of delay and variance in cellular doubling time (delay = 0%, standard deviation = 0 minutes), on network diameter and maximum edge degree by simulating the growth of exponentially dividing networks with a per division cell death probability ranging from 0.0-0.30. As predicted, mean network diameter monotonically increases with increasing cell death probabilities, increasing by ∼36% from a network diameter of 20 when cell death = 0.0 to 27.2 when cell death = 0.3 for networks of size 1,250, suggesting that cell death results in less densely branched networks (**Figure S12A**). Contrary to our expectations, the mean maximum edge degree peaks at intermediate values of apoptosis (0.10) rather than at apoptosis probability = 0.0. Beyond a cell death probability of 0.15, however, mean maximum edge degree monotonically decreases (mean maximum edge degree goes from 18.5 when cell death = 0.15 to 16.9 when cell death = 0.3 for networks of size 1,250), consistent with the idea that higher rates of apoptosis will lead to a reduction in maximum edge degree (**Figure S12B**). Taken together, these results begin to point to a potential alternative hypothesis for the evolution of elevated apoptosis rates in the snowflake yeast system: cell death may directly contribute to increased cluster size.

Next, we examined the joint effects of delay and apoptosis rate, varying delay from −50 to 50% of the second cellular doubling time, varying the cell death probability from 0.00 to 0.30, and holding the standard deviation constant (SD = 15 minutes). Increasing cell death probability increases network diameter regardless of the delay, and vice versa (**Figure S13A**), indicating no interaction between the two variables for this property. In contrast, the effect of increasing cell death probability on maximum edge degree depends strongly on the delay value. Specifically, when delay is high, increasing the cell death probability decreases maximum edge degree but increasing the cell death probability has little to no effect on maximum edge degree when delay is low (**Figure S13B**). The inverse, however, is not true; decreases in delay can always produce a decrease in maximum edge degree, but the effect is modulated by higher apoptosis rates (**Figure S13B**). Dynamic simulations of network growth and fragmentation show that size at fracture follows a similar trend: increasing cell death probability can lead to an increased size at fracture when delay is −40% or greater, but has no effect on size at fracture for clusters exhibiting a highly accelerated first division (**Figure S13C**). Simulations of fragmentation using the biophysical model showed qualitatively similar effects of delay and death probability, albeit with a significantly lower predicted size at fracture for the majority of parameter pairs; with increases (**Figure S14**; p < 0.05 after Bonferroni correction for all pairs except delay ≤ 25%, predictions becoming increasingly divergent as delay decreases or as cell death probability cell death = 0.00 & delay = 50%, cell death = 0.15, Wilcoxon test). Propagule proportion was the network property most profoundly influenced by apoptosis (**Figure S13D**). Increasing apoptosis quickly obscured the effect of delay and produced highly asymmetric fragmentation: at a cell death probability of 0.3, propagules contained only ∼10% of the cells of the parent cluster, while in the absence of apoptosis, propagules were ∼39% the size of their parents. Overall, while both delay and apoptosis rate modify network topology, delay has a larger and more consistent effect on size at fracture, whereas apoptosis has a much greater effect on the asymmetry of fragmentation.

## Discussion

Prior work on the snowflake yeast system identified several cellular traits (*e.g.*, increased cellular aspect ratio, increased cell size, and thicker and stronger cell-cell connections) that allow clusters to grow larger by changing the biophysical properties of the cells (23–25,28). Each of these, however, operates within the existing topology of the cluster, adjusting its mechanical or geometric properties without altering how cells are connected to one another. Here we identify two cellular traits, cell division timing and apoptosis rate, that exert control over multicellular morphology and size via topological changes that modify the network of cell-to-cell connections within a multicellular cluster rather than changing the biophysical properties of the constituent parts.

First, we found that asynchronous cell division caused by a first division delay produces more highly branched network topologies, enriched with “unbudded daughter” motifs, which concentrate crowding-induced mechanical stress on older cells and drive fragmentation at smaller sizes (**Figures 3, 4,** and **8**). However, not all asynchrony has this effect. Asynchrony driven by high variance in doubling time alone does not shift mean size at fracture, while asynchrony driven by an accelerated first division has the opposite effect of a delay, generating more filamentous branches and increasing size at fracture. To directly compare the relative contributions of biophysical and topological traits, we used the biophysical model implemented in **Figure S5** to estimate how much of the petite and grande cluster size difference is attributable to division synchrony, cell diameter, and cellular aspect ratio. Individually, these traits explained 35%, 26%, and −12% of the difference, respectively, but jointly explained 58% (**Table S3**). This indicates that, for our ancestral strains, division timing captures a substantial, largely independent share of the variation that biophysical traits alone cannot explain.

We then demonstrate how apoptosis rate modifies network topology, allowing clusters to reach a larger size by reducing the accumulation of localized crowding-induced mechanical stress around dead, non-dividing cells. While we framed cellular traits as acting on either a cluster’s biophysical properties or its topology, apoptosis shows that this distinction is not as clean as it first appears. Elevated rates of apoptosis in snowflake yeast were originally understood in purely biophysical terms: dead cells were hypothesized to act as weak points that facilitate fragmentation and thereby increase group-level fecundity (22,46,57). Our results add a complementary, topological mechanism: because a dead cell adds no further branches, apoptosis reshapes the network directly, increasing size at fracture while producing highly asymmetric fragmentation (**Figures S13**). The mechanical and topological consequences of cell death are therefore not mutually exclusive, and more broadly a trait’s biophysical and topological effects may be two intertwined consequences of the same cellular change. These consequences are also not fixed; apoptosis raises size at fracture for delayed and synchronous strains but has little effect when the first division is strongly accelerated (**Figure S13C**). The evolution of elevated rates of apoptosis, which has been observed repeatedly in snowflake yeast (22,56), also illustrates how the evolution of multicellularity opens up previously inaccessible paths to novel phenotypes; increased cell death would be strongly deleterious for a unicellular population of yeast grown under the same conditions. Whether a costly cell-level behavior pays off at the group level depends on the context created by other cell-level traits, an interaction that exists only at the level of the group. Because such benefits are expressed only in the multicellular context, these traits may also help entrench lineages in multicellularity (46,58,59), binding the fate of cells to that of the groups they build.

As the interaction between apoptosis and first division delay shows (Figure S13C), cellular traits do not act independently on group phenotype. Understanding how the cellular traits known to affect size at fracture interact with one another, including traits whose effects we have not yet characterized, is therefore an important next step, and future work should test whether similar interactions extend to bud site distribution and other traits shaping group phenotype. Bud site distribution is one such understudied trait, as we found that petite cells bud at wider angles than grande cells (47.4° vs. 35.2°), but what drives this difference, and what consequences it has for group-level properties, remains untested. In unicellular yeast, ploidy is known to affect bud site selection (60–62), and in multicellular yeast, altering the expression of *BUD4* can switch budding between axial and bipolar patterns, producing nearly a ten-fold difference in cluster size (47), making it another promising target for future work on the cellular determinants of group phenotype.

Our competition simulations (Figure 6) show that cell division timing is a cellular trait that has meaningful fitness consequences in a simple multicellular organism. The growth phase and settling selection components of our simulated competitions correspond closely to the two classical life history traits of fecundity and viability, respectively. Meanwhile, our manipulation of starting position during settling selection (random vs. top starting position) mirrors a shift from weak to strong selection on group-level phenotype driven by the environment. Viewed this way, our results show that synchronous division is favored under both weak and strong selection on group phenotype, and via both fecundity and viability-like fitness components, though its relative contribution to fitness depends heavily on the environment. This suggests that cellular traits capable of reshaping group topology, like division timing, can have fitness consequences even at this earliest stage of the transition to multicellularity.

These findings show how experimental evolution in a simple multicellular system, like the MuLTEE, can expose distinct mechanistic routes by which selection on cellular traits gives rise to novel multicellular phenotypes. Our results show how cellular traits affecting both biophysical and topological properties are plausible routes to controlling morphogenesis during the earliest stages of the transition to multicellularity. The relative importance of topological properties versus biophysical properties for different origins of multicellularity will likely depend on the specific cell biology of the unicellular ancestor and the mechanism of group formation. In multicellular groups formed via incomplete cell separation, intercellular bonds are often permanent, making topology a defining feature of their morphogenesis, whereas multicellular groups that form via aggregation tend to have reformable intercellular bonds (63). Thus, the same cell-level change could have different consequences for group-level phenotype depending on how the multicellular group is formed (accretive/clonal vs. aggregative group formation).

In extant multicellular organisms, the developmental programs that pattern the body are themselves multicellular adaptations (Bozdag et al., 2026), making it unclear how morphogenesis could first arise. For groups held together by permanent bonds, our results suggest how morphogenesis might get off the ground. When cells remain attached after division, division timing ceases to be a purely cellular property: it determines the topology of the group itself, affecting where mechanical stress accumulates and when the group fractures. Cell death acts in a similar way, because a branch stops developing when a cell stops dividing. Variations in cell division timing and apoptosis also shape how biomass is partitioned among offspring in multicellular groups. Neither process requires new gene products or cellular coordination, only quantitative shifts in the timing of cellular processes that unicellular ancestors already carry out. The raw material for this simple form of morphogenesis predates multicellularity itself. In this sense, cell division timing and apoptosis join previously described adaptations, such as increased cellular aspect ratio and stronger cell-cell connections, as candidate first steps toward morphogenetic control, with the difference that they act on topology rather than on biophysical properties. Taken together, our results show that two of the most basic cellular processes, when to divide and when to die, can shape the morphology and size of a nascent multicellular organism, providing a foundation upon which more elaborate, developmentally programmed patterning mechanisms might later evolve.

## Materials and Methods

### Soft-agar time-lapse preparation

The yeast strains were platted on YPD agar from 20% glycerol stock isolates stored at −80°C and grown for 3 days at 30°C in a static incubator. A single colony was picked and grown for 24 hours in 10 mL YPD in 25 × 150 mm culture tubes for 24 h at 30°C with 225 rpm shaking and then 1 mL sample was transferred to 10 mL of fresh media for a 4-hour growth for the strain to reach exponential growth phase. One milliliter of culture was centrifuged, the supernatant removed, and clusters were broken using a pipette tip. The sample was resuspended in 1 mL of 2x medium, subjected to 5 minutes of settling selection, and the top 100 μL collected. A final dilution was performed to achieve ∼200 clusters per 100 μL, based on microscopic estimation. Soft agar (0.7%) was prepared by melting in a microwave and allowed to cool slightly. Equal volumes (100 μL each) of cell suspension and soft agar were mixed and transferred to a microscopy plate well. The agar was allowed to solidify for at least 30 minutes before imaging. The time-lapses were recorded using a ×20 Nikon objective and images were collected every 5 minutes for 15 hours in total. Yeast growth during time-lapse microscopy occurred at room temperature (∼25°C). All yeast growth occurred at atmospheric oxygen levels. The same protocol was followed for all strains (ancestors and evolved isolates).

### Time-lapse analysis

Cell tracking and lineage reconstruction from time-lapse images was performed using Fiji (64), TrackMate (65), and Cellpose 2.0 (66). Manual correction was needed to ensure correct detection of the first frame the buds appeared to guarantee accurate measurement of cell doubling times. Then custom Python scripts were used to obtain the doubling time of the cells and the mother-daughter doubling time difference to quantify synchronous cell divisions. Cell diameter and cell aspect ratio were obtained from the last segmentation before the cell divides. The budding angle was calculated as the angle defined by the points of the mother’s cell distal pole, the center of the mother cell ellipse, and the proximal pole of the daughter cell (as a proxy for the bud scar). The proximal pole of each daughter cell was defined to be the pole closest to its mother cell’s center.

Some time-lapses were discarded before being processed if the yeast cells showed very little growth, if cells drifted out of the focal plane, or when there were too few usable doubling times because of crowding (from neighboring clusters or from within the same cluster). Another filtering step took place after the processing of the time-lapses was performed, the variance of the cellular doubling time distributions was checked via pairwise Fligner-Killen tests with Bonferroni correction for each of the time-lapses of the same replicate population and timepoint. Only the time-lapses that had a significant difference from the rest of the doubling time distributions were discarded, which led to the filtering of 3 time-lapses from 3 different evolved genotypes (PA3_t200_4_2022nov23, PA1_t1000_3_2022sep27, PA4_t200_2_2022nov23). For the grande ancestor, 8 time-lapses in total were analyzed from 5 different dates. Two of the dates (November 22^nd^, and 23^rd^ of 2022) show a lower doubling time that was not reproducible in the time-lapses performed later in 3 separate dates, therefore the time-lapses of those 2 original dates were discarded as they contained less cellular doubling times than the other time-lapses collected.

### Measuring cell properties from time-lapse microscopy

To confirm that cell properties were maintained during the time-lapses, cell growth was calculated from the time-lapse data. This information was obtained by processing the spots and edges files output from TrackMate’s time-lapse analysis to link all masks along each cell’s growth trajectory and counting the number of divisions by classifying mother and daughter cells after each cell division. From this dataset, the second division of the ancestors was used to calculate the median cell diameter (long-axis; TrackMate reports the semi-axis length, so the reported values were multiplied by 2) and aspect ratio for each cell. The second division was chosen because the median of the cell properties from the masks in the second division showed less variation than the final mask of the first cell division, which could be affected by errors in cell segmentation due mainly to cell overlap. The median cell properties from all cells were then used to calculate the overall median cell diameter and aspect ratio per strain for comparison with the static measurements.

### Measuring cluster and cell properties from static images

Ancestral strains were grown in a YPD agar plate and 5 colonies of each were grown in different tubes of liquid YPD for 24 hours. After the first growth cycle a random sample of 100 µL was transferred to a new tube for another day to grow, and from the remanent of the tube after day 1 and with the tube of day 2, the measurements were performed.

For the cluster size measurements, a sample of 1 mL of the culture was taken, and then two dilutions were performed in series in water, a 1:100 and a 1:10 for a 1:1000 total dilution. Then 500 µL was transferred to 16 well-plate for imaging of the whole well using a 4x Nikon objective. The images were analyzed with an ImageJ script to calculate the cluster properties.

To measure the cell shapes, the cell wall was stained using calcofluor. For this process a sample of 100 µL was collected from each tube, and it was washed in PBS two times, and then it was incubated in 1 mL PBS with 1 µL of calcofluor and incubated for 10 minutes in the dark and washed one last time before being imaged with a 40x Nikon objective. The images were processed with Cellpose 2.0 (66) to create the masks of the cells and manual correction was performed to delete the masks of all cells that were not fully grown or haven’t budded yet, and after the masks and images were processed in ImageJ to obtain the measurements of the cells. Finally, to obtain the distribution of cells in the cluster of the aerobic ancestor, the same cells stained with calcofluor where imaged using a 20x Nikon objective by adding 3 µL into the microscope slide and gently flattening them into a monolayer of cells. Enough images were taken to capture at least 100 clusters per replicate. To measure the number of cells per cluster, cell segmentation was performed with cellpose2 to segment all the cells, and the cluster segmentation and assignment of cell masks into a cluster was performed with a custom Fiji script.

### Log-normal distribution parameterization

To create the log-normal distributions we defined the parameters for a normal distribution by selecting the mean (*μ*) and standard deviation (*σ*), and the parameters for the log-normal distribution were obtained by the formulas:

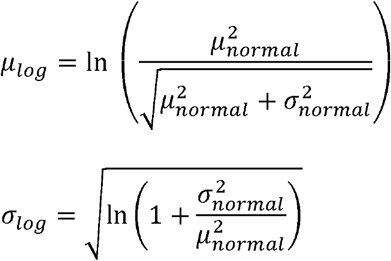

In addition, to simulate the higher variation observed in the first division when a delay was observed, the standard deviation of the first division was scaled by the formula:

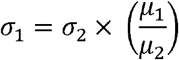

Where *μ*_1_ and *σ*_1_ correspond to the mean and standard deviation of first cellular division, and *μ*_2_ and *σ*_2_ are the mean and standard deviation of subsequent cellular divisions.

### Network model (using discrete or synthetic continuous data)

The complete network model was implemented in Python 3.10 using the NetworkX library (67). Clusters are represented as a dynamical network graph G = (N, E), in which cells are nodes (N) and incomplete cell divisions as edges (E). Cell divisions are modeled by adding a new node connected to the nodes representing the respective mother cell. Upon creation, the new node and the mother node were attributed a division time which was sampled from empirical data (Figures 3, 4, 6, and S3D, S5-S9) or from a log-normal distribution (Figures 5, 8, S3A-C, S11, and S12-S14). After addition of all new cells, the network was evaluated for potential fragmentation events, which were determined by using our created metric the edge degree:

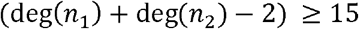

where deg(*n*_1_) and deg(*n*_2_) are the node degrees of the nodes forming an edge. Edge degree was evaluated for every node n in the network. If more than 1 edge was flagged for fragmentation, one was selected at random.

Some code used GNU Parallel (68) to coordinate workflow of scripts and to increase the execution speed.

### Normalized network diameter estimation

The normalization of the network diameter was used to compare the network topology of the simulated networks to an exponentially growing network independent of the size (Figure S6). To find the formula for the normalized network diameter, an exponential network was grown, and a linear regression was fit to the data of the log scaled number of nodes and network diameter, and from the regression parameters the formula was estimated to be:

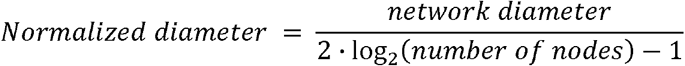

### Settling selection simulations

To mimic competition experiments between different strains, we first simulated the growth phase of the cluster followed by calculating the sedimentation velocities in order to perform settling selection. For the growth phase, clusters were sampled from an initial cluster size distribution and initiated in a 1:1 ratio of both competing strains. These initial clusters were grown for 24h or until the carrying capacity of the simulation is reached (carrying capacity is 10 million cells unless specified otherwise).

Next, the settling selection is performed consisting first of a 10% dilution of the total biomass (total number of cells), and then the selection of the first 10% of clusters that reach the end of the length of a tube (2.5 cm), when starting the cells at a random position or at the top of the tube, and the surviving cells are going to be transferred to the next growth phase for the defined amount of cycles or until one population goes extinct. The growth phases were performed in Python using the network model described above, and the settling simulations were performed using a spatial model in MATLAB to calculate the diameter of the clusters (24). To estimate which clusters will reach the bottom of the tube and survive during settling selection, we calculate the cluster diameter from the simulated clusters and use Stokes’ law to estimate the velocity of sedimentation (27,53). Each cluster diameter is obtained by calculating the radius of center of the cluster, and then transformed to the sphere diameter using the formula 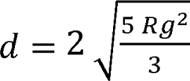 gyration, obtained by calculating the root-mean-square distance between the cells and the where *Rg* is Radius of gyration (69). The sedimentation velocity is calculated as:

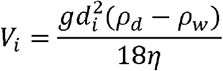

where *d_i_* is the diameter of the cluster, *g* is the gravity constant (*g* = 9.81 *ms*^—2^), *ρ_d_* is the mass density of yeast cells (*ρ_d_* = 1112.6 *kg m*^3^; Bryan et al., 2010), *ρ_w_* is the mass density of water at 25°C (*ρ_w_* = 997 *kg m*^3^), and *η* is the dynamic viscosity of water at 25°C (*η* = 9.81 × 10^—4^ *pa s*).

### Biophysical model

In this biophysical model, cells are represented as ellipsoids with defined diameter and aspect ratio parameters. New cells are added by sampling both the budding location and growth angle, if the sampled position is too close to an already present budded cell a new position is going to be sampled, there is a maximum of 10 attempts to find a location for the new cell that is not so crowded, but if the position can’t be found then the last sampled position is going to be used. Mechanical stress is going to be estimated as the overlap between cells, therefore cluster fragmentation occurs when the overlap between cells meets or exceeds the specified threshold. In the cases where the clusters are built from networks (Figure 6 and Figs. S5, S11 and S14), the cells are added in order in which the nodes were added to the network.

### Cluster size difference predictions using a spatial model

The goal of this procedure is to approximate the experimental cluster size measurements with the size at fracture predicted by the spatial MATLAB model (24), specifically examining three key variables: cell aspect ratio, cell diameter, and cell synchrony. For each individual variable, we performed a controlled analysis where we first calculated the overlap threshold needed to trigger fragmentation at the target volume while holding all three variables constant to the grande strain values. After establishing this threshold, we then calculated the predicted fracture size using the specific parameters of each ancestral strain (petite and grande) for the variable being tested, while keeping the other two variables constant at reference values. The contribution of each variable to the observed size difference between ancestors was quantified as the percentage of the experimental difference explained by the simulated difference. For both cell aspect ratio and cell diameter analyses, we implemented synchronous cell division as the standard condition for both strains.

## Statistical tests

All statistical tests were performed using R version 4.1.2 (2021-11-01) (71). For consistency, we chose to use non-parametric tests throughout the manuscript because some distributions violated the normality assumption. Wilcoxon tests, paired with Cliff’s delta (δ) for effect size, were performed to test the significance in the difference between median values of all distributions analyzed in this manuscript, including from empirical or simulation sources. Fligner-Killeen test was used to test the difference in variance between the doubling time distributions of the ancestors doubling time data. Wilcoxon signed-rank test, paired with correlation coefficient (r) for effect size, was performed to test if there was a significant difference in median doubling time between mother and daughter cells, which corresponds to paired data. A Bonferroni correction was used for multiple test correction in the cases where multiple or pairwise tests were performed.

For the evaluation of the “bustiness” of cell divisions (**Figure S3**), event times were binned into 5-minute windows, matching the temporal resolution of the empirical time-lapse data, to generate a time series of division counts per window for each simulated run. We used an autocorrelation function (ACF) to estimate a “synchrony index” which measures how similar a time series is to a time-lagged copy of itself, as a function of the lag interval. A population dividing synchronously is expected to show high autocorrelation at a lag close to one mean doubling time, whereas a desynchronized population shows little autocorrelation at that time. Because of the exponential growth nature of the simulation, each bin was normalized by dividing each bin counts by the population size present in the population at the start of that bin to keep the synchrony signal in the cell division timing. For each run, the reported ACF value was the highest autocorrelation found within a window spanning 0.5–1.5 times the population’s expected mean doubling time. This yielded one ACF value per simulated run, which was then compared across conditions. For the variance and delay sweeps (ordered, multi-level conditions), a Spearman’s rank correlation test was used to test for a monotonic relationship between condition level and ACF value, since this test does not assume normality or a linear relationship. For the two-group comparison between grande and petite ancestor cells, a Wilcoxon rank-sum test was used instead.

All R scripts are included in the GitHub repository associated with this manuscript.

### Use ofLLMs in writing and coding

LLM assistance was used for some aspects of writing and coding of the scripts used to generate the results presented in this manuscript. For writing, Claude (Sonnet 3.5 and newer) was used to suggest style edits for early versions of the manuscript. All AI-suggested edits were carefully reviewed to guarantee that the content of the edited text was preserved and that the resulting text still accurately represented the authors’ intent. For later versions of the manuscript, all modifications were performed without any AI assistance. For coding, Claude (Sonnet 3.5 and newer) was used to aid the development of the following: creating bash scripts including the ones used for parallel execution of scripts and the workflow of the settling simulations; modifications to the MATLAB script for the biophysical simulations; modifications to the base network simulations; and refinement of the figures presented in this article. AI assistance was not used in the creation of the base network simulation functions or the Python and ImageJ scripts used to analyze the microscopy data. All AI-suggested additions to code were carefully reviewed and tested to verify that the new code worked as intended.

## Data availability

Raw images from time-lapse and static microscopy can be accessed at: https://doi.org/10.6019/S-BIAD4067. Processed image data, raw data from simulations, and all code used for simulations and analyses is available at: https://github.com/pipecedeno/cell_division_timing_shapes_the_morphology_and_size_of_nascent_multicellular_organisms.

## Acknowledgments

We thank Peter J. Yunker, Tom E.R. Belpaire, and G. Ozan Bozdag for their thorough comments on the manuscript. This work was supported by NIH grant No. 5R35GM138030 to W.C.R. The Multicellularity Long-Term Evolution Experiment is supported by NSF LTREB grant DEB-2452109.

## Author Contributions

**Conceptualization**: Luis F. Cedeño-Pérez, William C. Ratcliff, Peter L. Conlin

**Data Curation:** Luis F. Cedeño-Pérez

**Formal Analysis**: Luis F. Cedeño-Pérez

**Funding Acquisition**: William C. Ratcliff

**Investigation**: Luis F. Cedeño-Pérez

**Methodology**: Luis F. Cedeño-Pérez, Peter L. Conlin

**Project Administration:** William C. Ratcliff, Peter L. Conlin

**Software**: Luis F. Cedeño-Pérez, Peter L. Conlin (network model implementation and simulation pipelines); Rozenn M. Pineau (code for settling selection simulation); Thomas C. Day (biophysical simulation and synchronous bursting code)

**Supervision:** William C. Ratcliff, Peter L. Conlin

**Validation:** Luis F. Cedeño-Pérez, William C. Ratcliff, Peter L. Conlin

**Visualization**: Luis F. Cedeño-Pérez, Thomas C. Day, Peter L. Conlin

**Writing – Original Draft Preparation**: Luis F. Cedeño-Pérez, Peter L. Conlin

**Writing – Review & Editing**: Luis F. Cedeño-Pérez, Rozenn M. Pineau, Thomas C. Day, William C. Ratcliff, Peter L. Conlin

## Supplementary Materials

Supplementary Figures 1-14

Supplementary Tables 1-3

Supplementary Videos 1-2

## Notes

### Competing Interest Statement

The authors have declared no competing interest.

### Summary of Updates

New analysis and figures has been added, including the characterization of synchronous bursting in the ancestral strains, the effects of apoptosis in fragmentation, and the comparison between both network and biophysical models. In addition, some parts of the main text have been updated, including parts of the introduction, results, and discussion.

