## Supplemental Files for "Cell division timing shapes the morphology and size of nascent multicellular organisms"

**Supplementary Figures**

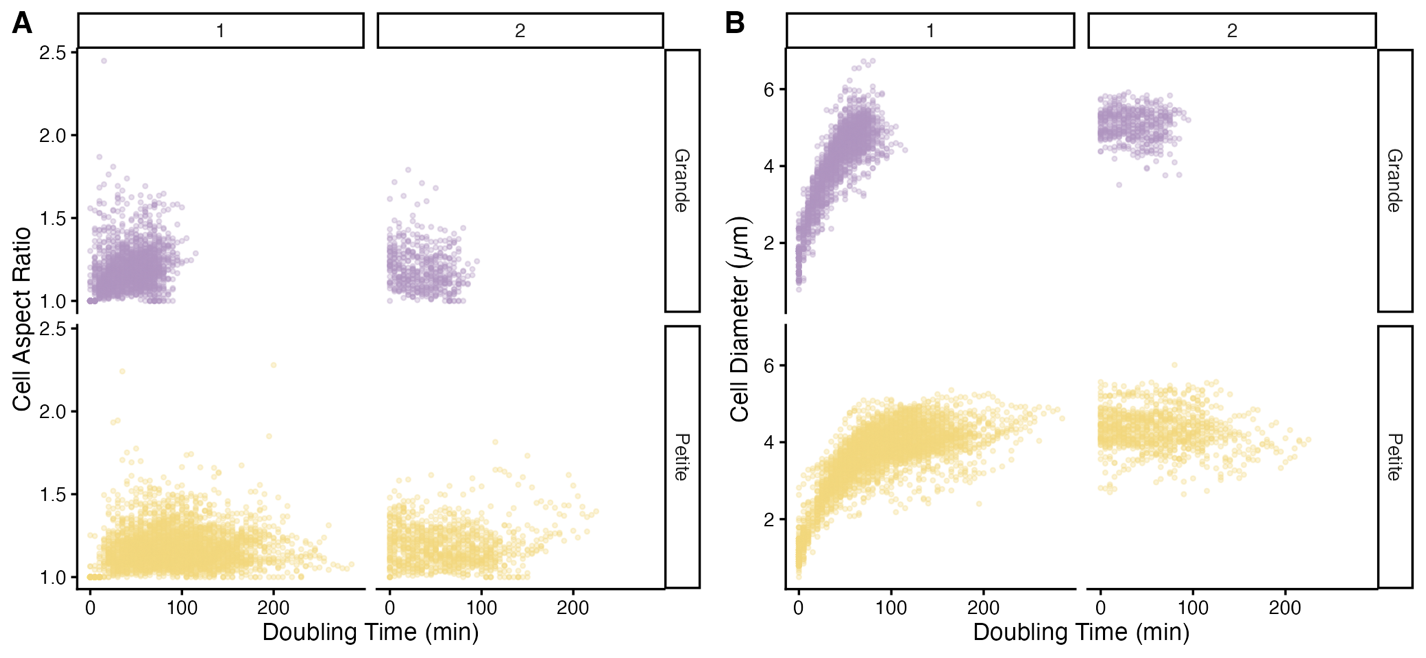

**Figure S1. Measurement of cellular properties during time-lapse experiments.** **A**) Cell aspect ratio changes during the first and second cell division for the petite and grande ancestors for all cell doublings measured in time-lapses. **B**) Changes in cell size as measured by cell diameter during the first and second cell division. All time-lapse videos were recorded using a ×20 Nikon objective and images were collected every 5 minutes for 15 hours in total.

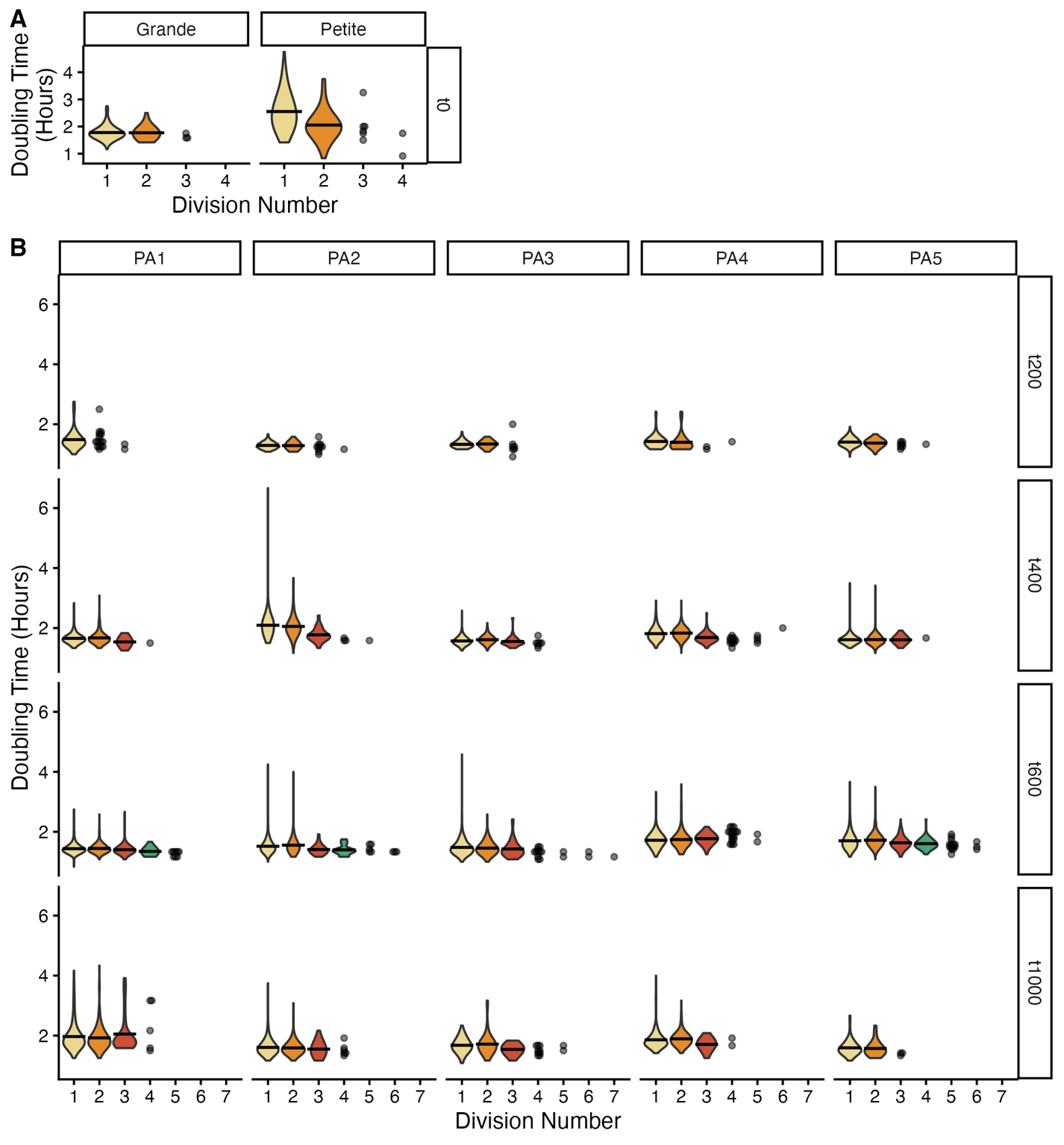

**Figure S2. Consistency in mean doubling times across multiple cellular divisions.** Mean doubling times are shown for successive divisions in the 600-day evolved anaerobic strains, demonstrating that after the first division, subsequent divisions maintain similar mean timing, data shown for the ancestral strains (**A**) and evolved petite strains (**B**). Only divisions with n ≥ 20 observations are shown in violin plots, cell divisions with less than 20 data points are shown directly. This pattern supports the key assumption used in our models that doubling times remain consistent after the first division.

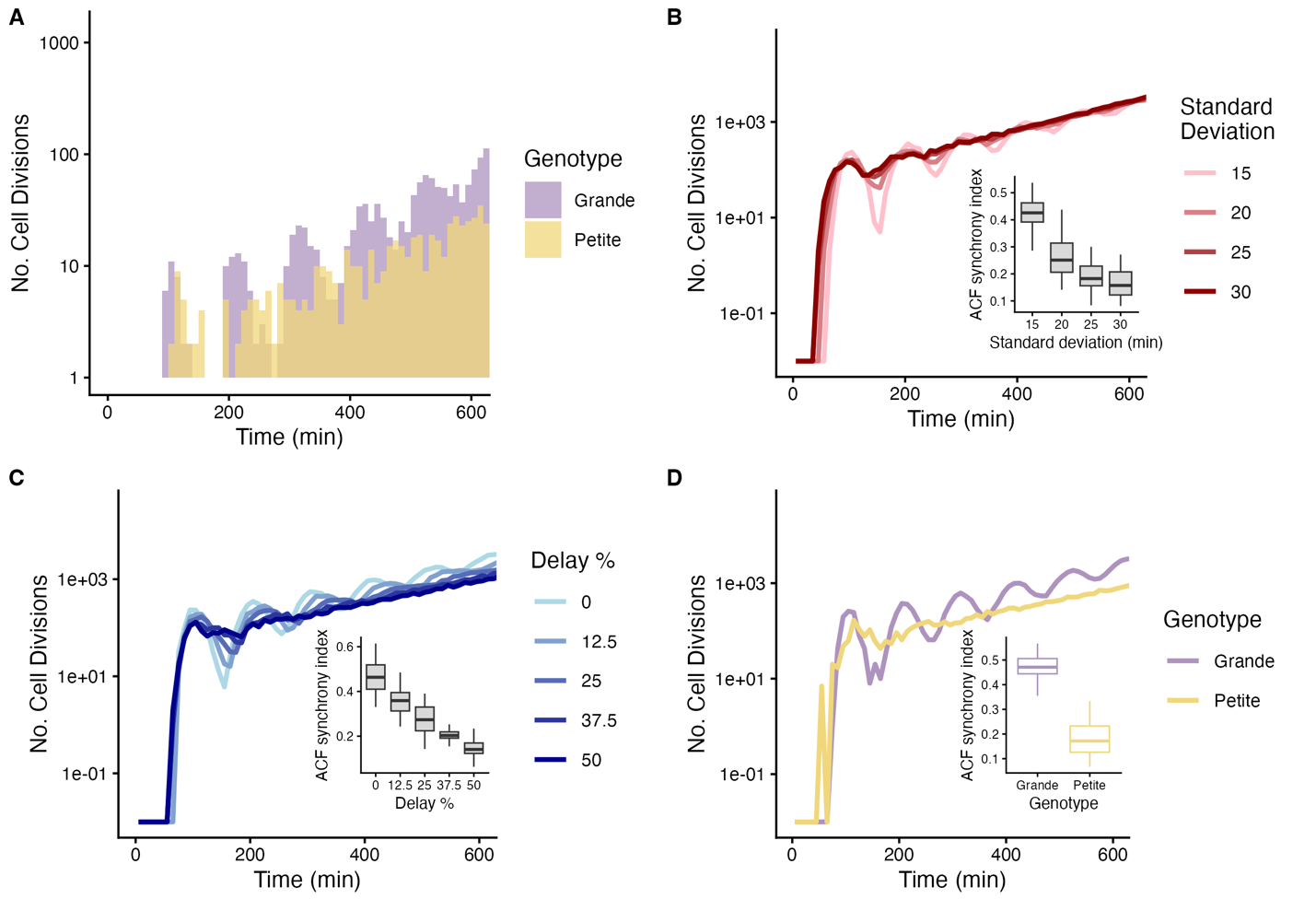

**Figure S3. First division delay and increased variance in cellular doubling time can eliminate synchronous bursting in cell division timing.** **A**) Example simulation of cellular doubling events generated by sampling the empirical doubling times measured for the grande and petite ancestor. **B-C**) Effects of increased variance and first division delay in cell division synchronous bursting. Autocorrelation function (ACF) synchrony index (insets) declined with increasing standard deviation of the doubling-time distribution (ρ = −0.78, *p* < 2.2x10^-16^, Spearman's rank correlation coefficient). Similar effect was observed by increasing first division delay (ρ = −0.88, *p* < 2.2x10^-16^, Spearman's rank correlation coefficient). **D**) Grande and petite cellular doubling events generated by sampling with replacement the empirical doubling time distributions. Grande ancestor maintains a higher level of synchrony compared to the petite ancestor (ACF synchrony index shown in inset of 0.48 compared to 0.18, respectively; *p* < 2.2x10^-16^, Wilcoxon test). All cell doubling counts are binned in windows of 10 minutes for visualization, and 5 minutes for statistical analysis (same time scale of empirical data), 30 simulations were performed for each parameter combination shown in panels B, C and D starting from 30 cells (15 mother and 15 daughter cells) for a total simulation time of 1 thousand minutes.

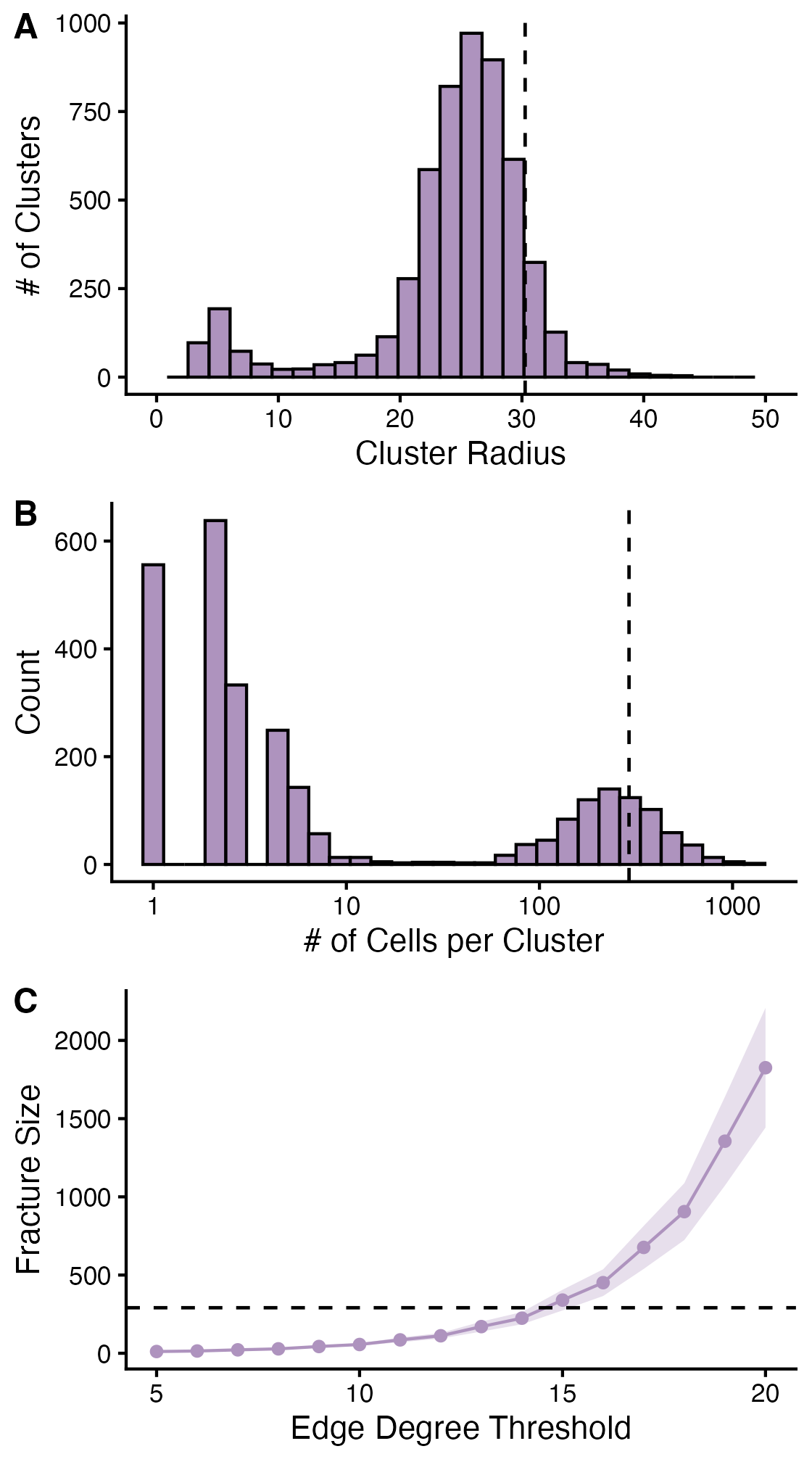

**Figure S4. Edge degree threshold calibration based on grande ancestor fracture size.** **A**) Cluster size distribution of the aerobic ancestor after ~10 generations (2 days of growth). Dashed vertical line represents the 90th percentile, corresponding approximately to the 30 μm cluster radius at expected fragmentation (28). **B**) Distribution of cells per cluster in the aerobic ancestor. Dashed vertical line represents the 90th percentile (291 cells per cluster). **C**) Mean fracture size distributions for different edge degree thresholds using aerobic strain doubling time distributions. An edge degree threshold of 15 produces clusters that fracture at 339 cells, closely approximating the target size of 291 cells per cluster (dashed horizontal line). Shaded area represents standard deviation. 300 simulations were performed for each edge degree threshold.

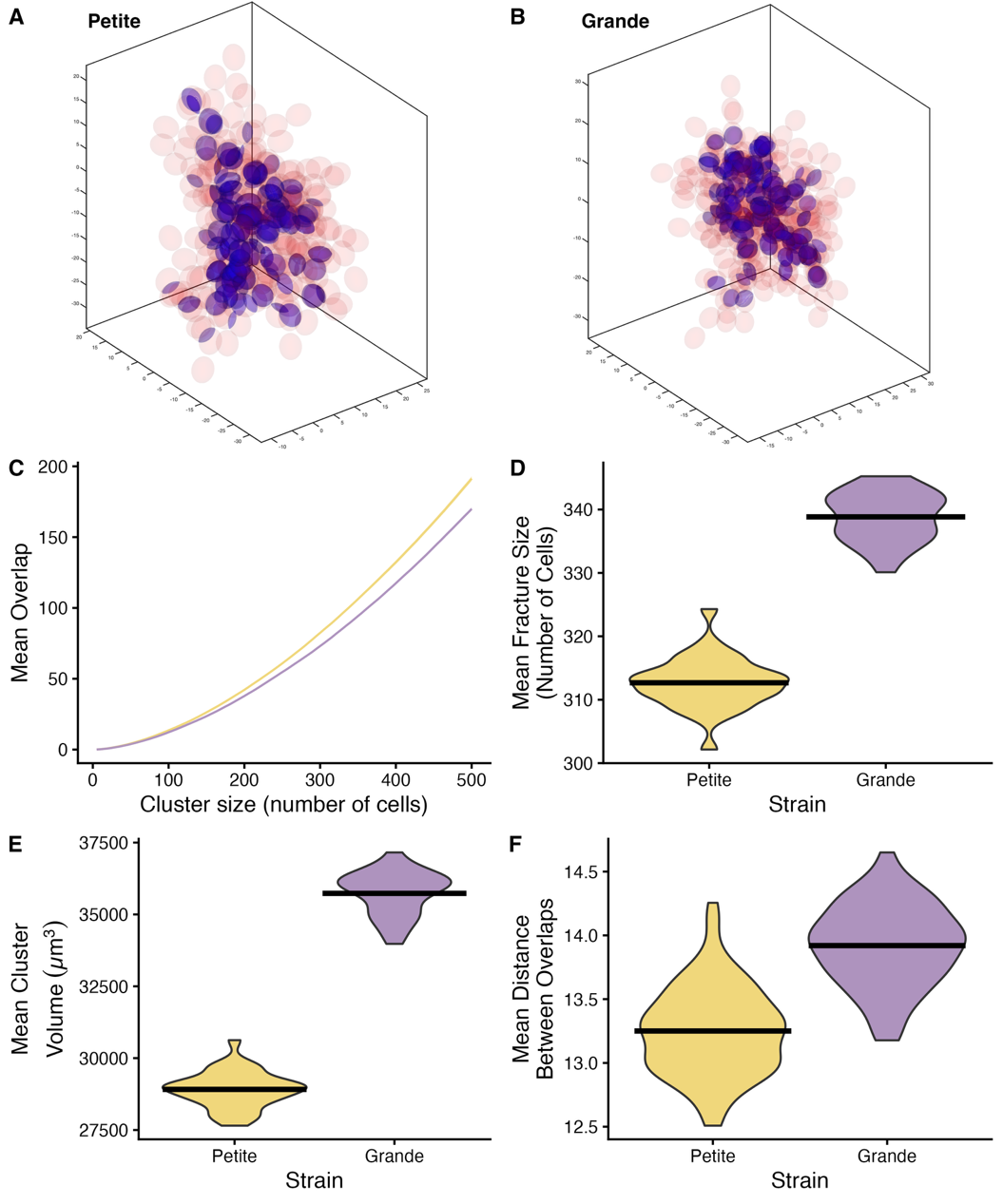

**Figure S5. Asynchronous division promotes localized mechanical stress accumulation and smaller cluster size.** **A**-**B**) Representative clusters of petite (**A**, 302 cells) and grande (**B**, 329 cells) ancestors at fragmentation, with cells shown in red and regions of cell-cell overlap in blue. **C**) Accumulation of cell-cell overlaps during cluster growth in petite (yellow) and grande (purple) ancestors. 100 clusters were analyzed, and for each cluster 30 simulations were performed, so the line represents the mean of the means of the cluster simulations, and the shaded area shows the confidence interval of the population mean. **D**) Distribution of cluster sizes at fragmentation for petite (yellow, median 312.7 cells) and grande (purple, median 339.3 cells) ancestors (*p* < 2.2x10^-16^, Wilcoxon test). **E**) Mean cluster volume distribution at fracture for petite and grande ancestors (median volume 28,976 μm³ and 35,983 μm³, respectively; *p* < 2.2x10^-16^, Wilcoxon test). **F**) Distribution of mean distances between overlap centers. For each cluster, distances were calculated between 10,000 randomly sampled overlap pairs. Petite networks show significantly shorter mean distances between overlaps (median distance between overlaps: petite = 13.26 μm, grande = 13.92 μm, *p* < 2.2×10^-16^, Wilcoxon test), confirming more localized mechanical stress. For **D**, **E** and **F**, data represent means across 30 simulations for each of 100 independent clusters.

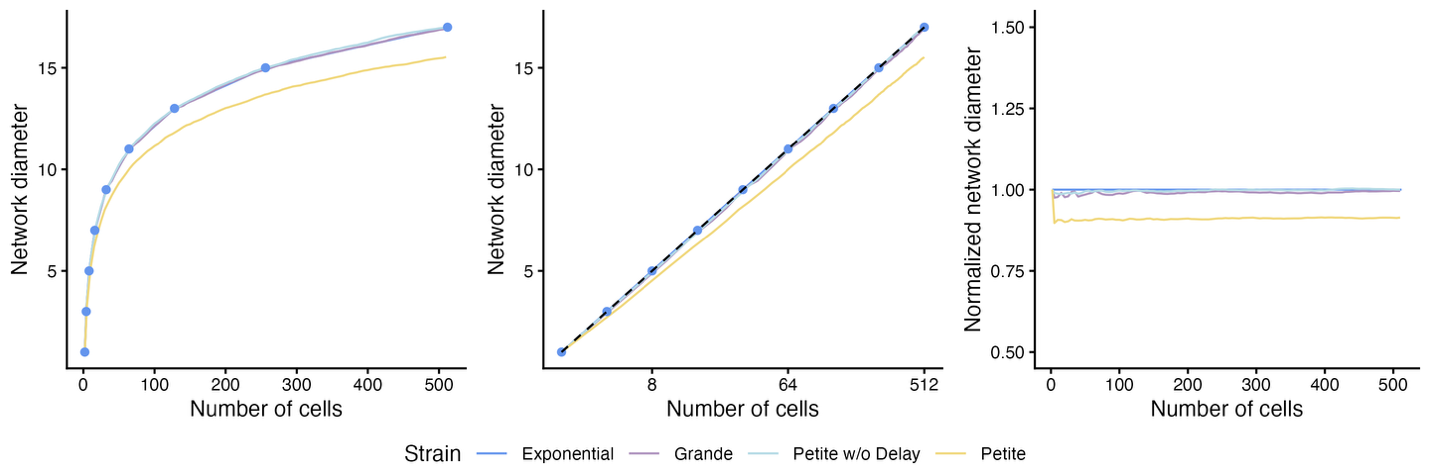

**Figure S6. Normalization method for network diameter. A**) Relationship between cell number and network diameter in exponentially growing networks, showing that diameter increases with cluster size. **B**) Log-transformation of cell number reveals a linear relationship with network diameter that can be accurately predicted using linear regression. **C**) Network diameter values normalized to exponentially growing networks of equivalent size, where a value of 1 represents the expected diameter for a perfectly exponential network. This normalization approach enables comparison of network topologies across different cluster sizes, allowing quantification of whether a network is more or less branched compared to an exponentially growing network with synchronous divisions. See Methods for the normalization equation.

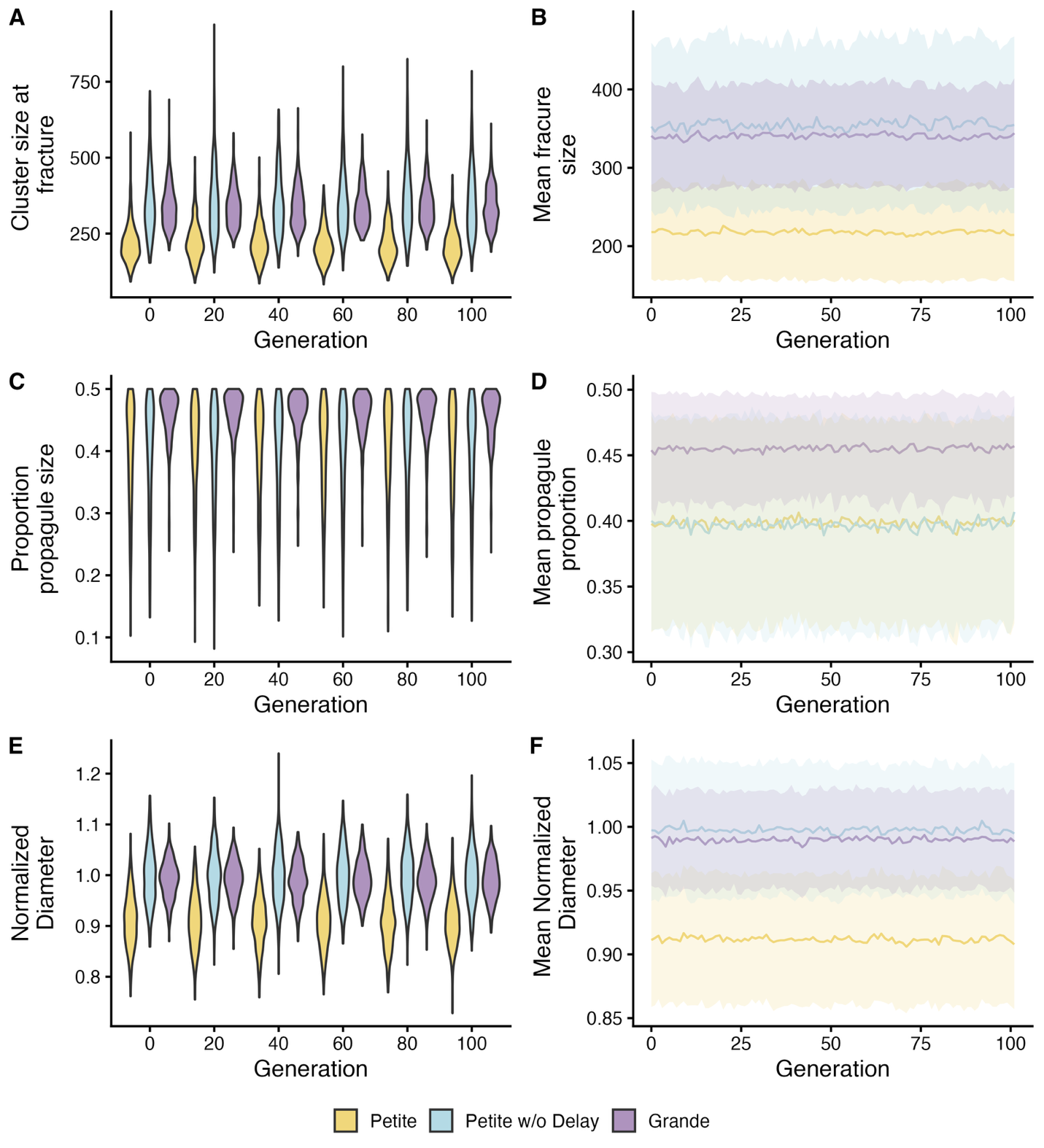

**Figure S7. Stability of network properties over 100 cluster generations.** Simulations tracked either the parent or propagule cluster, randomly selected for continued growth in each generation. **A**, **C**, **E**) Distributions of network properties shown at 20-generation intervals. **B**, **D**, **F**) Mean and standard deviation of network properties for each generation. All plots show no change in the shape of the distributions or in its mean value.

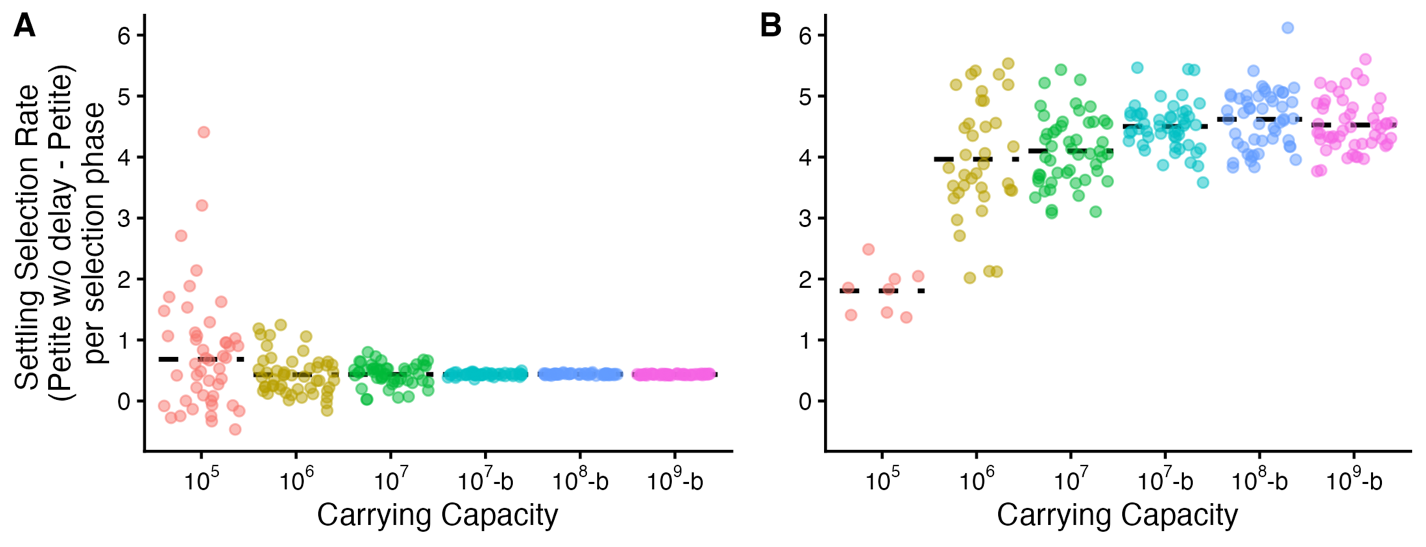

**Figure S8. Impact of carrying capacity on settling selection rate. A**) Settling selection rates with clusters starting at random positions. **B**) Selection rates with clusters starting at the top. Bootstrap simulations (marked in the plot by the “-b” after the carrying capacity) involved initial growth to 1 million cells, followed by cluster diameter calculation and resampling to 10% of the new carrying capacity as if already performing the first dilution of the selection regime. No further growth phases were simulated post-settling selection. Pairwise Wilcoxon tests showed no significant differences in means across carrying capacities for **A** after Bonferroni correction (*p* > 0.05). In **B**, the 10^5^, 10^6^ and 10^7^ carrying capacities were statistically different than the rest (*p* < 0.005, pairwise Wilcoxon tests with Bonferroni correction). Each dot represents the mean selection rate per simulation for each of the phases, in total 50 simulations were performed for up to 20 transfers. For the 10 thousand cells simulation in **B**, less than 50 data points are shown because one of the populations went extinct in the first settling selection and therefore the selection rate couldn’t be calculated.

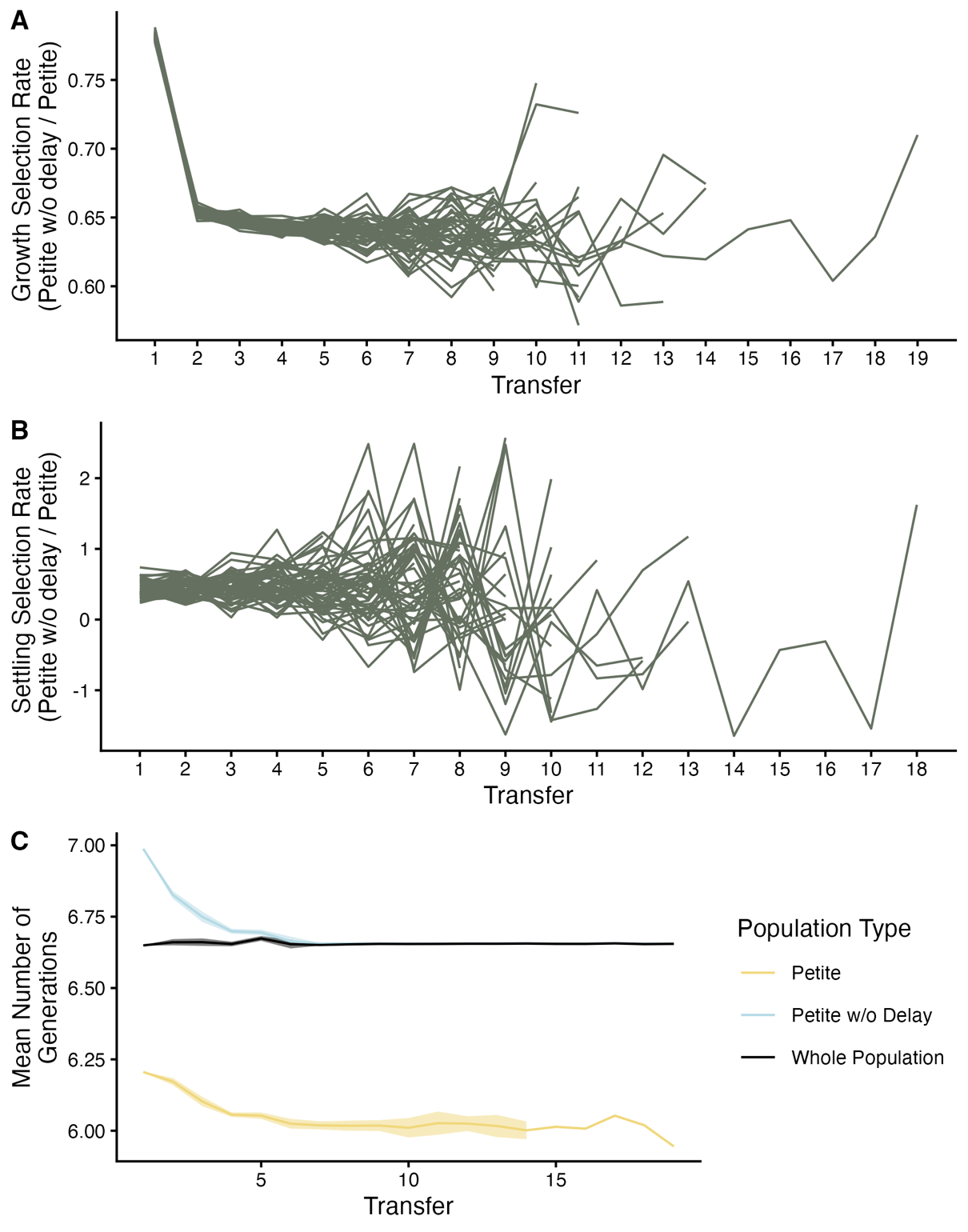

**Figure S9. Differential generation count between fast and slow strains drives enhanced initial growth selection rate.** **A**) Growth selection rate, showing a notably higher value for the first transfer compared to subsequent transfers. **B**) Settling selection rate, does not exhibit the elevated first-transfer phenomenon observed in the growth phase. **C**) Mean number of generations achieved by each strain and the total population per transfer. The maximum difference in growth between strains occurs during the first transfer, explaining the elevated initial growth selection rate. Shaded areas represent one standard deviation from the mean. Lines in panels **A** and **B** represent selection rates from independent simulations (n = 50 simulations).

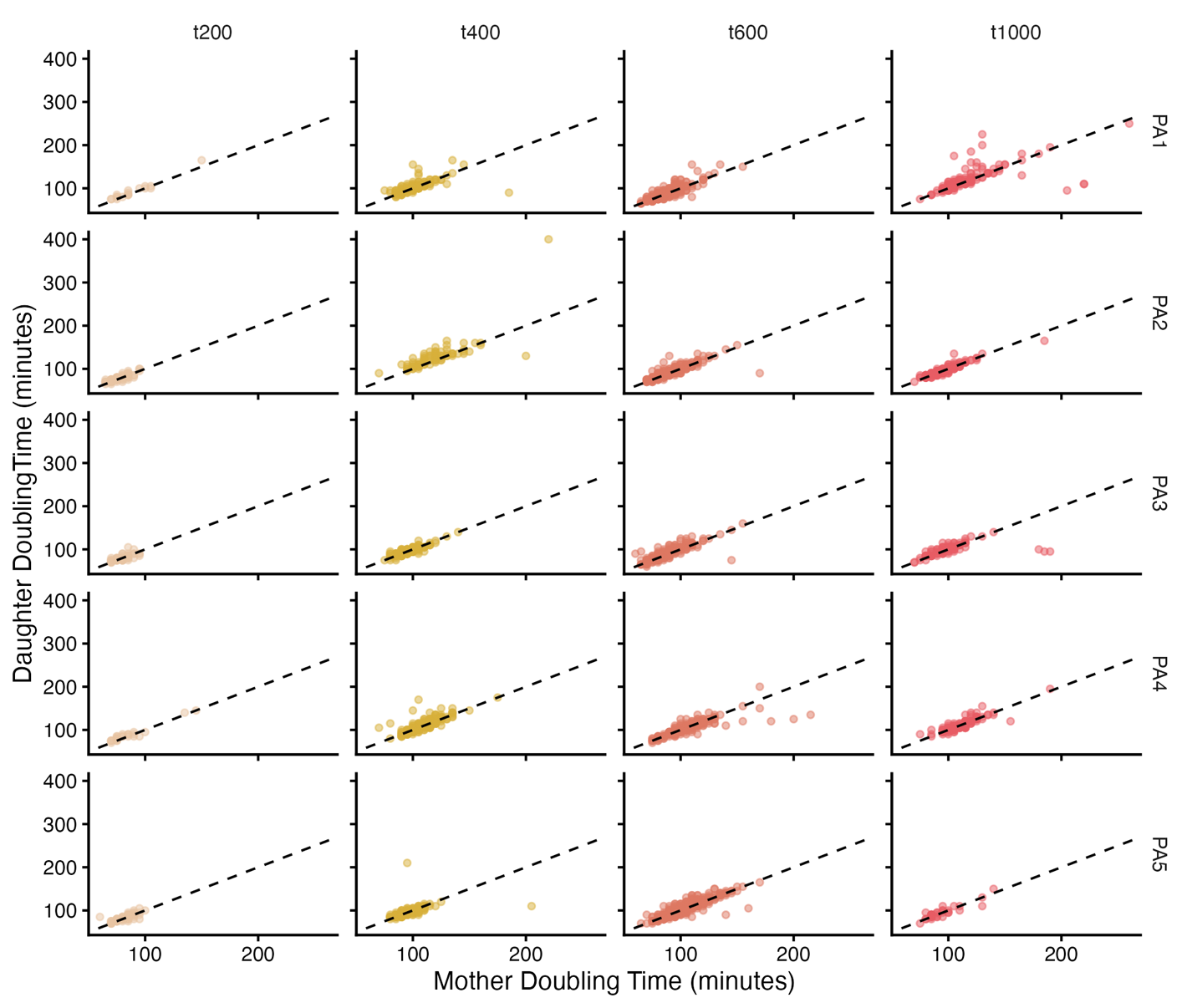

**Figure S10.** **All evolved strains from the petite ancestor show high levels of synchrony between mother and daughter cell divisions.** Scatter plots of mother vs. daughter cell doubling times. The identity line (dashed line) means that both mother and daughter cells are dividing at the same time. The following strains have a slower daughter cell division compared to the mother cell (*p* < 0.0025 with Bonferroni correction for 6 out of 20 strains, Wilcoxon rank-sum test): PA1 t400, and t1000; PA2 t400, and t600; PA3 t600; and PA5 t600.

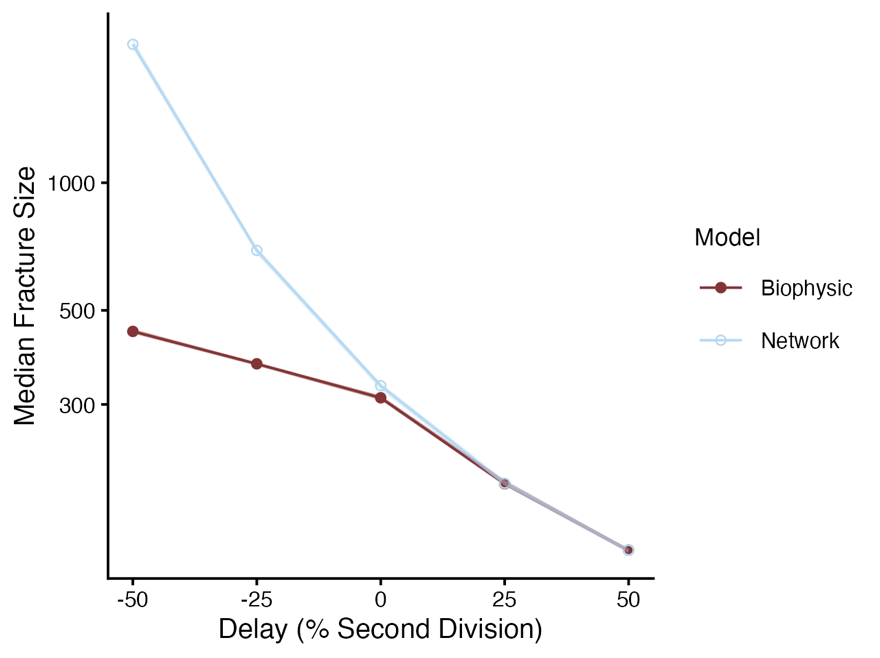

**Figure S11. Divergence in predicted size at fracture for biophysical and network models across a range of delay values.** The biophysical and network model predict the same median size at fracture for first division delays of 25% and 50% (*p* > 0.01 after Bonferroni correction, Wilcoxon test). Meanwhile, the biophysical model predicts a lower size at fracture than the network model for synchronous or accelerated first division (delay values of 0%, -25%, and -50%, *p* < 0.01 after Bonferroni correction, Wilcoxon test). However, the biophysical model still predicts an increase in fragmentation size when comparing the accelerated first divisions to the synchronous cell divisions (*p* < 2x10^-16^, pairwise Wilcoxon test). These results were obtained by first using the network model to simulate 20 cluster fragmentations and randomly selecting either the parent or propagule after fragmentation. The networks were then loaded into the biophysical simulation, adding cells in the same order in which the nodes were added in the network simulation, until fragmentation was triggered by the overlap accumulation between cells. Each point depicts the median fracture size for 30 separate trials at each delay value across all 20 observed fragmentation events (n = 600 per combination).

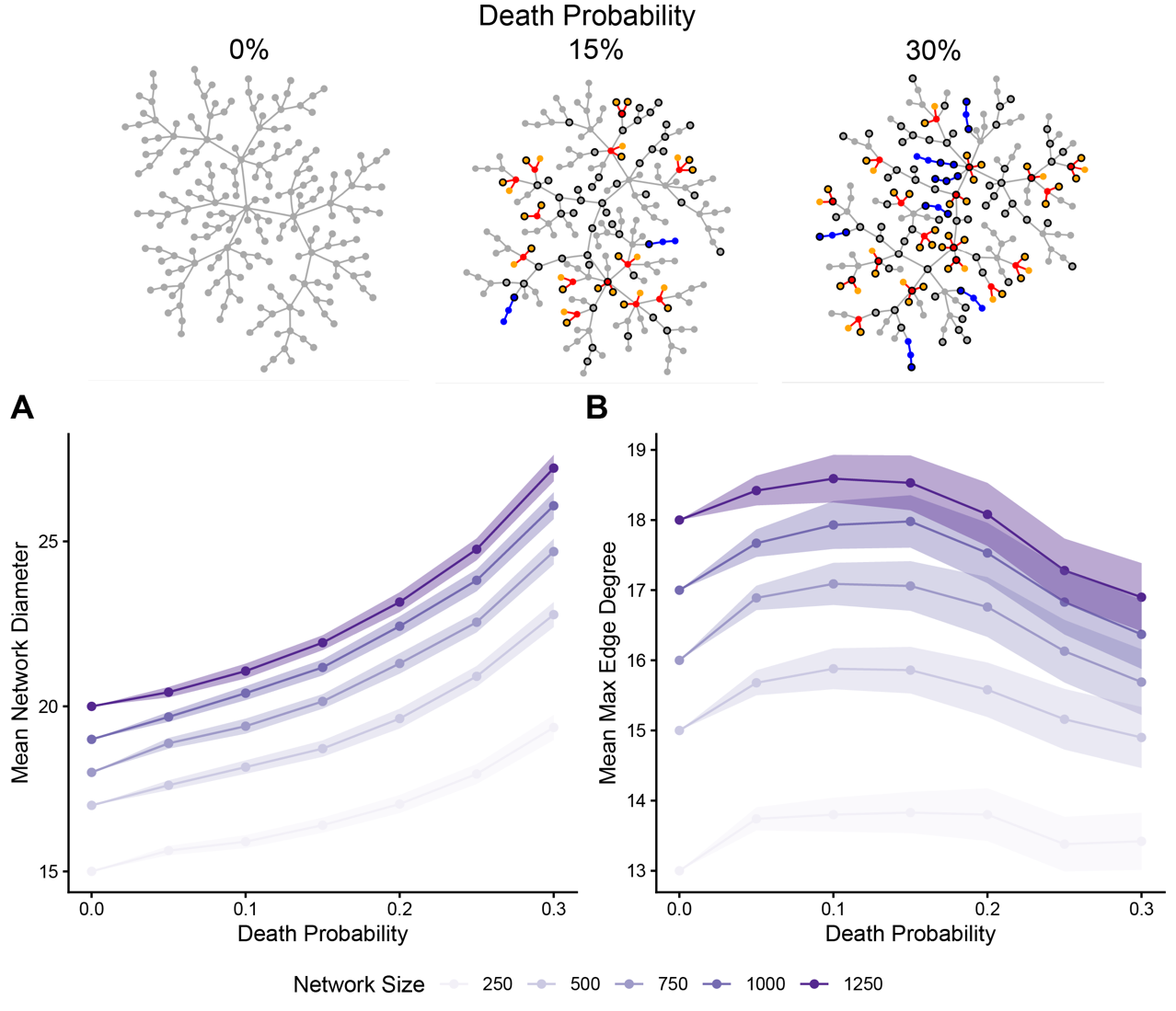

**Figure S12. Cell death increases network size by reducing the accumulation of maximum edge degree**. **A**) Relationship between network diameter and death probability. **B**) Relationship between mean maximum edge degree and death probability. Both metrics demonstrate that increasing death probability reduces network crowding, supporting the idea that increased cell death leads to an increase in fragmentation size. For each of seven death probabilities (0 to 0.3 in 0.05 increments), 100 independent simulations were performed, with networks grown from single nodes to 1300 nodes. Cell death was modeled as a constant probability that a cell stops dividing after each cell division. Nodes with black borders represent cells that died during growth. We illustrate the effects on cluster topology with example networks of 200 nodes at different levels of cell death (Top).

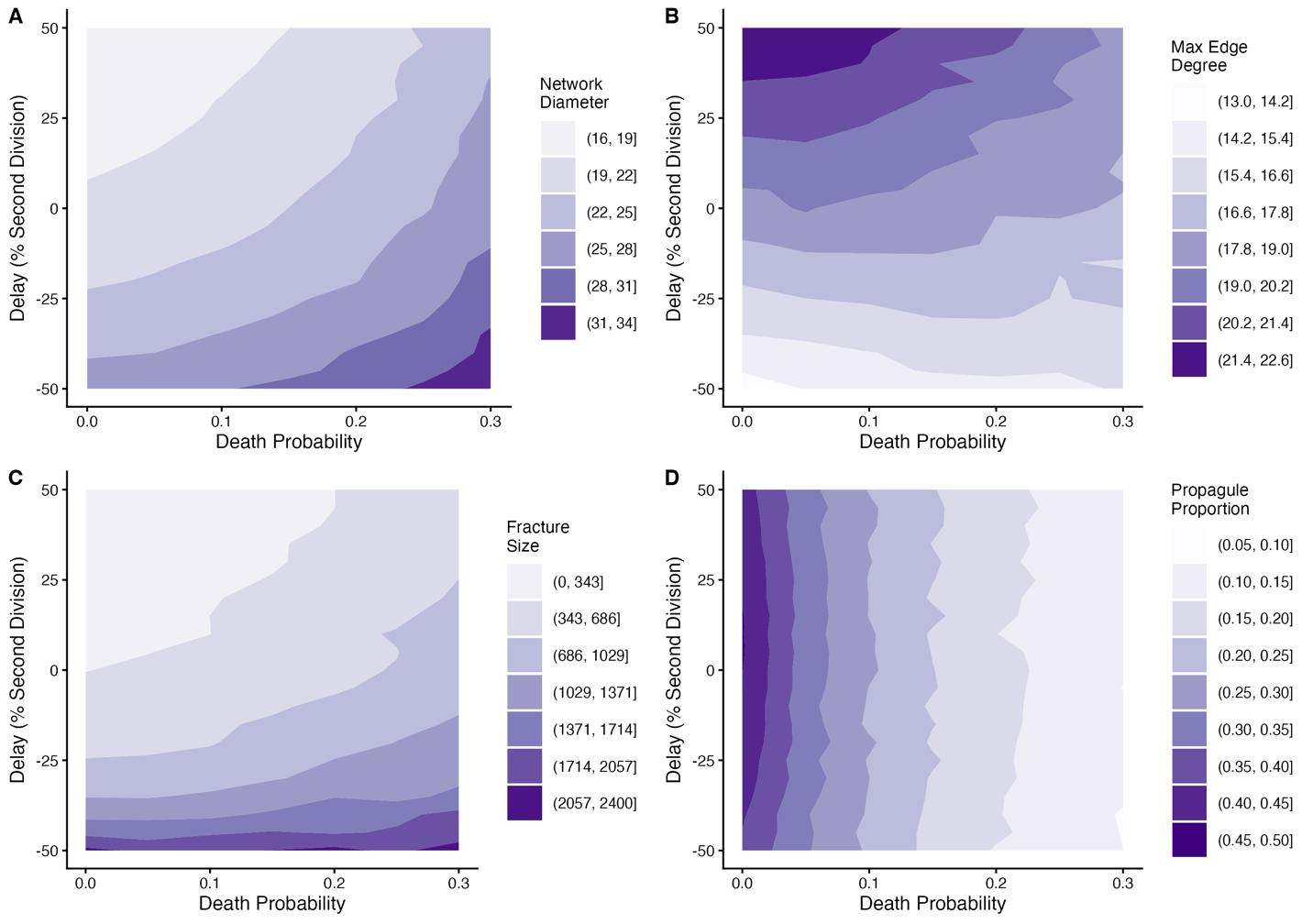

**Figure S13. Combined effects of delay and cell death probability on network properties and multicellular fragmentation.** Panels showing the effect of death probability and first division delay on the network variables: network diameter (**A**), maximum edge degree (**B**), size at fracture (**C**), and propagule proportion (**D**). Delay and cell death probability exhibit no interaction of network diameter. Cell death probability has a modulating effect on delay for maximum edge degree, as higher death probabilities reduce the effect on maximum edge degree caused by delay. Cell death probability increases fragmentation size when an accelerated first division is absent. Cell death probability drastically changes the topology of the networks causing a highly asymmetric fragmentation. For panels **A** and **B,** 100 independent simulations were performed, with networks grown from single nodes to 1300 nodes. Both panels show diameter and maximum edge degree values for networks at 1300 nodes. For panels **C** and **D**, 30 simulations with fragmentation were performed for 50 cluster generations growing the parent or propagule at random. The values displayed are grand means: the mean of the 30 simulations was calculated for each cluster generation, and these 50 cluster generation means were then averaged (n = 50 per delay and death probability combination). All the simulations performed used the combination of seven cell death probabilities (0 to 0.3 in 0.05 increments) and 21 delay values.

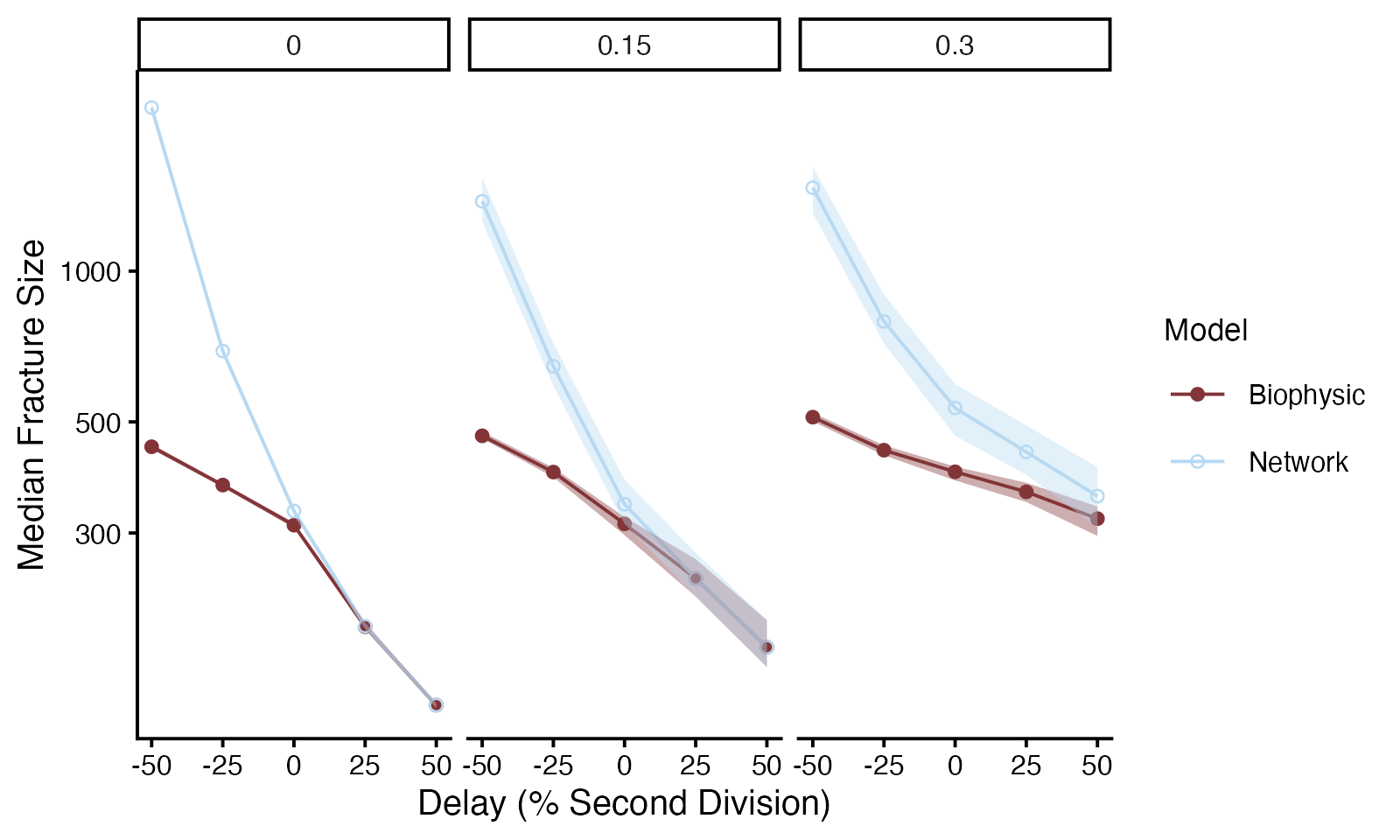

**Figure S14. Divergence in predicted size at fracture for biophysical and network models across a range of delay values and cell death probabilities.** Both network and biophysical models show that delay is a major predictor of fracture size and that apoptosis causes higher fragmentation sizes for delayed and synchronous strains. However, the predicted size at fracture is significantly lower in the biophysical model for the majority of parameter pairs (*p* < 0.05 after Bonferroni correction for all pairs except delay $\leq$ 25%, cell death = 0.00 & delay = 50%, cell death = 0.15, Wilcoxon test). Simulations were performed as in **Figure S11**, with clusters constructed by adding the cells in the same order as in the network simulations into the biophysical framework for 20 cluster fragmentations for 30 different trials. Each point depicts the median fracture size for 30 separate trials at combination of delay and cell death values across all 20 observed fragmentation events (n = 600 per combination).

**Table S1. Summary of time-lapse observations and cell division data.** The table provides the total number of time-lapse experiments and observed cell doublings, as well as a breakdown of doubling events by division number for the petite and grande ancestors.

| Division number | Number of cell doublings (petite ancestor) | Number of cell doublings (grande ancestor) |
| --- | --- | --- |
| 1 | 135 | 144 |
| 2 | 48 | 36 |
| 3 | 6 | 3 |
| 4 | 2 | 0 |
| Total | 191 | 183 |

**Table S2. Summary of time-lapse measurements of each replicate population in the anaerobic line.** The table contains both the amount of time-lapses collected and the number of cell doublings measured.

| Strain | T200 time-lapses | T200 cell doublings | T400 time-lapses | T400 cell doublings | T600 time-lapses | T600 cell doublings | T1000 time-lapses | T1000 cell doublings |
| --- | --- | --- | --- | --- | --- | --- | --- | --- |
| PA1 | 2 | 70 | 6 | 635 | 11 | 1801 | 6 | 358 |
| PA2 | 3 | 257 | 4 | 279 | 8 | 930 | 4 | 448 |
| PA3 | 2 | 136 | 8 | 641 | 5 | 802 | 4 | 274 |
| PA4 | 2 | 92 | 10 | 867 | 6 | 662 | 4 | 392 |
| PA5 | 4 | 242 | 6 | 535 | 14 | 2111 | 2 | 104 |

**Table S3. Contribution of cellular traits to cluster size differences between grande and petite strains at fracture.** The predictions were made using a previously validated biophysical model.

|  | Grande Volume ($\mu m^{3}$) | Petite Volume ($\mu m^{3}$) | Volume Difference (Grande - Petite) ($\mu m^{3}$) |
| --- | --- | --- | --- |
| Experimental data | 98,756 | 55,187 | 43,569 |
| **Predictions** |  |  |  |
| Cell aspect ratio | 68,579 | 73,919 | -5,339 (-12%) |
| Cell diameter | 113,355 | 101,987 | 11,368 (26%) |
| Cell synchrony | 105,374 | 90,199 | 15,175 (35%) |
| All combined | 105,374 | 79,924 | 25,449 (58%) |
